# Regenerative failure of sympathetic axons contributes to deficits in functional recovery after nerve injury

**DOI:** 10.1101/2025.01.08.631956

**Authors:** Tina Tian, David Kim, Kuai Yu, Criss Hartzell, Patricia J. Ward

**Affiliations:** Medical Scientist Training Program, Emory University School of Medicine, Atlanta, GA, USA 30307; Neuroscience Graduate Program, Laney Graduate School, Emory University, Atlanta, GA, USA 30307; Department of Cell Biology, Emory University School of Medicine, Atlanta, GA, USA 30307

## Abstract

Renewed scientific interest in sympathetic modulation of muscle and neuromuscular junctions has spurred a flurry of new discoveries with major implications for motor diseases. However, the role sympathetic axons play in the persistent dysfunction that occurs after nerve injuries remains to be explored. Peripheral nerve injuries are common and lead to motor, sensory, and autonomic deficits that result in lifelong disabilities. Given the importance of sympathetic signaling in muscle metabolic health and maintaining bodily homeostasis, it is imperative to understand the regenerative capacity of sympathetic axons after injury. Therefore, we tested sympathetic axon regeneration and functional reinnervation of skin and muscle, both acute and long-term, using a battery of anatomical, pharmacological, chemogenetic, cell culture, analytical chemistry, and electrophysiological techniques. We employed several established growth-enhancing interventions, including electrical stimulation and conditioning lesion, as well as an innovative tool called bioluminescent optogenetics. Our results indicate that sympathetic regeneration is not enhanced by any of these treatments and may even be detrimental to sympathetic regeneration. Despite the complete return of motor reinnervation after sciatic nerve injury, gastrocnemius muscle atrophy and deficits in muscle cellular energy charge, as measured by relative ATP, ADP, and AMP concentrations, persisted long after injury, even with electrical stimulation. We suggest that these long-term deficits in muscle energy charge and atrophy are related to the deficiency in sympathetic axon regeneration. New studies are needed to better understand the mechanisms underlying sympathetic regeneration to develop therapeutics that can enhance the regeneration of all axon types.

## INTRODUCTION

The sympathetic nervous system is responsible for a myriad of homeostatic mechanisms ranging from control of vascular tone and bronchodilation to blood glucose and thermoregulation (McCulloch et al., 1967; Grassi, 1998; Dibona, 2002; Carnagarin et al., 2018; Gagnon and Crandall, 2018). Recently, it has been shown that postganglionic sympathetic axons are important for maintaining neuromuscular junction (NMJ) integrity and health via postsynaptic β2-adrenergic receptors (ADRB2s) (Calvey et al., 2015; Khan et al., 2016; Senger et al., 2018; Sayanagi et al., 2021). Furthermore, sympathetic signaling at the NMJ through ADRB2s has been demonstrated to be important for skeletal muscle mitochondrial protein synthesis and respiration (Robinson et al., 2011; Khan et al., 2016; Azevedo Voltarelli et al., 2021). Despite the broad range of sympathetic nervous system functions and the fact that sympathetic axons comprise nearly a quarter of the axons in the rat sciatic nerve (Schmalbruch, 1986; Tian et al., 2024), they remain an understudied class of neurons in the context of peripheral nerve injuries (PNIs).

PNIs are common and are a significant public health issue with few patients achieving full functional recovery (Scholz et al., 2009); however, the current standard of care still relies on the slow and inefficient process of spontaneous regeneration with few patients achieving full functional recovery (Rodríguez et al., 2004). Over the last few decades, perioperative electrical stimulation (ES) has been shown to enhance motor and sensory axon regeneration (Gordon et al., 2007; Asensio-Pinilla et al., 2009; Gordon et al., 2009; Gordon et al., 2010; Huang et al., 2013; Calvey et al., 2015; Elzinga et al., 2015; Gordon, 2016; Gordon and English, 2016; Willand et al., 2016; Borschel, 2019; Halevi et al., 2019; Park et al., 2019; Power et al., 2020; Zuo et al., 2020a; Zuo et al., 2020b; Javeed et al., 2021; Sayanagi et al., 2021; ElAbd et al., 2022; Keane et al., 2022; Roh et al., 2022), but the effects of ES or other neuronal stimulation interventions on sympathetic regeneration are largely unknown. ES promotes axon regeneration via the upregulation of neurotrophins and regeneration-associated genes, which are similar to mechanisms seen in conditioning lesions (CLs) (McQuarrie et al., 1977; Forman et al., 1981; McQuarrie and Grafstein, 1981; McQuarrie, 1986; Hoffman, 2010; Senger et al., 2018; Chan, 2019b; Senger et al., 2019). The CL assay involves injuring the nerve prior to a test lesion which leads to enhanced motor and sensory regeneration (McQuarrie et al., 1977; McQuarrie et al., 1978; Forman et al., 1981; McQuarrie and Grafstein, 1981; Bisby and Pollock, 1983; McQuarrie, 1986); however, its clinical use is limited. Notably, although CLs are mechanistically similar to ES, they have been demonstrated to inhibit sympathetic axon regeneration (McQuarrie et al., 1978). With ES now available as a treatment for patient populations (Al-Majed et al., 2000; Al-Majed et al., 2004; Gordon et al., 2010; Wong et al., 2015; Barber et al., 2018; Chan, 2019a; Piccinini et al., 2020; Power et al., 2020; Borschel, 2023), there is a need to characterize how it may affect sympathetic axon regeneration and functional recovery. Furthermore, the newly appreciated functions of sympathetic innervation in muscle may play important roles in achieving full functional recovery after PNIs.

We hypothesized that the lumbar sympathetic neurons respond to growth-promoting interventions with enhanced axon regeneration, similar to their motor and sensory counterparts. To test our hypothesis, we compared motor and sensory axon regeneration to sympathetic regeneration in the acute setting in response to ES, CL, and a temporally controlled, neuronal-specific approach called bioluminescent optogenetics (BL-OG). BL-OG was employed both *in vivo and in vitro*. We also evaluated long-term sympathetic axon reinnervation of skin and muscle, both anatomically and functionally, in response to ES. Our findings suggest that lumbar sympathetic neurons do not follow the same principles we have long appreciated regarding motor and sensory axon regeneration in response to classic regeneration-enhancing interventions.

## MATERIALS AND METHODS

### Animals

All experiments were conducted on approximately equal numbers of males and female adult (8-25 weeks old) mice weighing 15-32 g. All experiments were approved by the Institutional Animal Care and Use Committee of Emory University (PROTO202100177).

Mice were randomly assigned to No Treatment (transection only), CL (a treatment not dependent on the electrical properties of the axons), or ES groups to calculate regenerative indices in the sciatic nerve in the acute regeneration assay. For the long-term regeneration assays, mice were randomly assigned to sham ES (Sham) or ES groups. Tg(Thy1-YFP)16Jrs transgenic mice (Jackson laboratory stock no.003782, referred to as “YFP-16”) were used for the aforementioned assays. The YFP- 16 mouse model was chosen to allow for differential visualization of motor and sensory axons versus sympathetic axons. The YFP-16 mouse exhibits robust expression of yellow fluorescent protein (YFP) in somatic motor neurons and large-caliber somatic sensory neurons but exhibits negligible YFP expression in postganglionic sympathetic axons **(Supplemental Figure 1)** (Feng et al., 2000; Taylor-Clark et al., 2015), which can then be subsequently visualized via immunohistochemistry (IHC). A separate cohort of C57BL/6J (Jackson laboratory stock no. 000664, referred to as “wild type” or “WT”) was used for retrograde tracing experiments after No Treatment, CL, or ES because the presence of a fluorescent reporter was not needed.

Acute sympathetic regeneration, *in vivo* and *in vitro*, was also assessed using BL-OG to allow for a more specific method of stimulation of the postganglionic sympathetic neurons (Gomez-Ramirez et al., 2020; Crespo et al., 2021; English et al., 2021; Ecanow et al., 2022; Ikefuama et al., 2022). The current clinical paradigm for ES is unlikely to elicit action potentials in the small-caliber unmyelinated sympathetic neurons due to the reversal of the size-recruitment principle with ES (Blair and Erlanger, 1933; Fang and Mortimer, 1991; Singh et al., 2000; Llewellyn et al., 2010). Furthermore, ES and CL non-selectively affect many types of neurons alongside surrounding non-neuronal cells, such as macrophages and Schwann cells (Huang et al., 2010; Kearns and Thompson, 2015; McLean and Verge, 2016; Gu et al., 2022), so we used BL-OG as a more specific method of stimulation. Th(Th-cre)Fl12Gsat/Mmucd transgenic mice (MMRRC stock no. 0.17262-UCD, referred to as “ThCre”) were crossed with R26-LSL- LMO3 transgenic mice (Jackson laboratory stock no. 034853, referred to as “eLMO3”) to create the ThCre:eLMO3 mouse model. Mice with either the ThCre or eLMO3 construct, but not both, are not responsive to coelenterazine (CTZ), a substrate to the luciferase enzyme which is conjugated to the eLMO3 luminopsin, as both constructs together are needed for Cre-inducible BL-OG-mediated chemogenetics (Crespo et al., 2021; English et al., 2021; Medendorp et al., 2021). Therefore, mice expressing either one of the singular constructs were used as controls. Final numbers of mice in each experimental group are listed in **Table 1**. **Figure 1** presents a summary of methods for the acute regeneration assays while **Figure 7** presents a summary of methods for the long-term regeneration assays.

**Figure 1:**
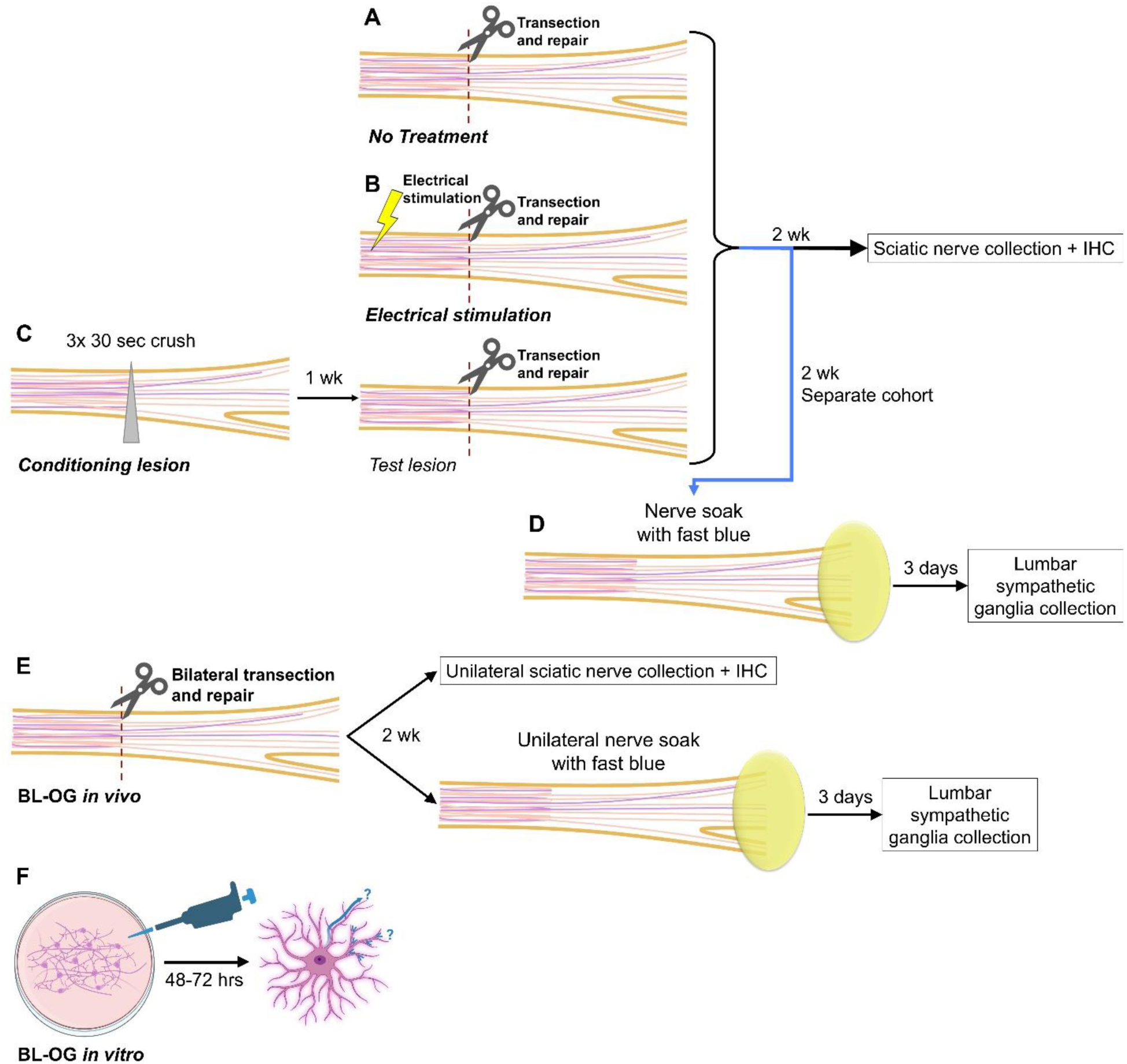
Acute regeneration summary of methods. Evaluation of sympathetic regeneration following (A) No Treatment, (B) electrical stimulation, or (C) conditioning lesion with immunohistochemistry (IHC) and (D) fast blue retrograde tracing. (E) Evaluation of sympathetic regeneration following bioluminescent optogenetics (BL-OG) *in vivo*. (F) *In vitro* evaluation of the effects of BL-OG on sympathetic regeneration by evaluating the neurite elongation (enclosing radius) and neurite branching (branching index). Partially created with BioRender.com.

**Table 1:** Experimental groups. Outline of the number of animals used in each experiment. The last column represents the final number included in the analyses. *1-4 animals were excluded due to poor quality nerve sections (distal stump did not reach 3000 μm or the transection site was not visible), incomplete extraction of the lumbar sympathetic ganglia, poor tissue quality, inadequate electrode placement, or unexpected death. **Cohort randomly selected from long-term cohort in the row above.

FIGURES AND LEGENDS
| Experiment | Experimental Group | Genotype | Sex |
| --- | --- | --- | --- |
| 2-wk regeneration after ES and CL | No Treatment | YFP-16 | 3 M, 3 F* |
|  | ES | YFP-16 | 4 M, 4 F |
|  | CL | YFP-16 | 4 M, 4 F |
| 2-wk retrograde tracing after ES and CL | No Treatment | WT | 3 M, 3 F |
|  | ES | WT | 3 M, 3 F |
|  | CL | WT | 3 M, 3 F |
| Bioluminescence quantification | Control | eLMO3 | 2 M, 2 F |
|  | BL-OG | ThCre:eLMO3 | 2 M, 2 F |
| Extracellular field recordings | Control | eLMO3 | 3 M, 1 F |
|  | BL-OG | ThCre:eLMO3 | 2 M, 1 F |
| BL-OG *in vivo* sciatic nerve regeneration | Control | ThCre | 4 M, 5 F* |
|  | BL-OG | ThCre:eLMO3 | 4 M, 3 F* |
| BL-OG *in vivo* retrograde tracing | Control | ThCre | 6 M, 4 F* |
|  | BL-OG | ThCre:eLMO3 | 5 M, 3 F* |
| BL-OG *in vitro* regeneration | Control | eLMO3 | 2 M, 2 F |
|  | BL-OG | ThCre:eLMO3 | 2 M, 2 F |
| Long-term functional recovery via sweat and purine quantification | Sham | YFP-16 | 3 M, 4 F* |
|  | ES | YFP-16 | 3 M, 4 F* |
| Long-term NMJ receptor quantification** | Intact | YFP-16 | 3 M, 2 F |
|  | Sham | YFP-16 | 2 M, 2 F |
|  | ES | YFP-16 | 2 M, 2 F |
| Long-term functional recovery via EMG and muscle mass | Sham | YFP-16 | 4 M, 5 F* |
|  | ES | YFP-16 | 5 M, 4 F* |

**Table 2:**
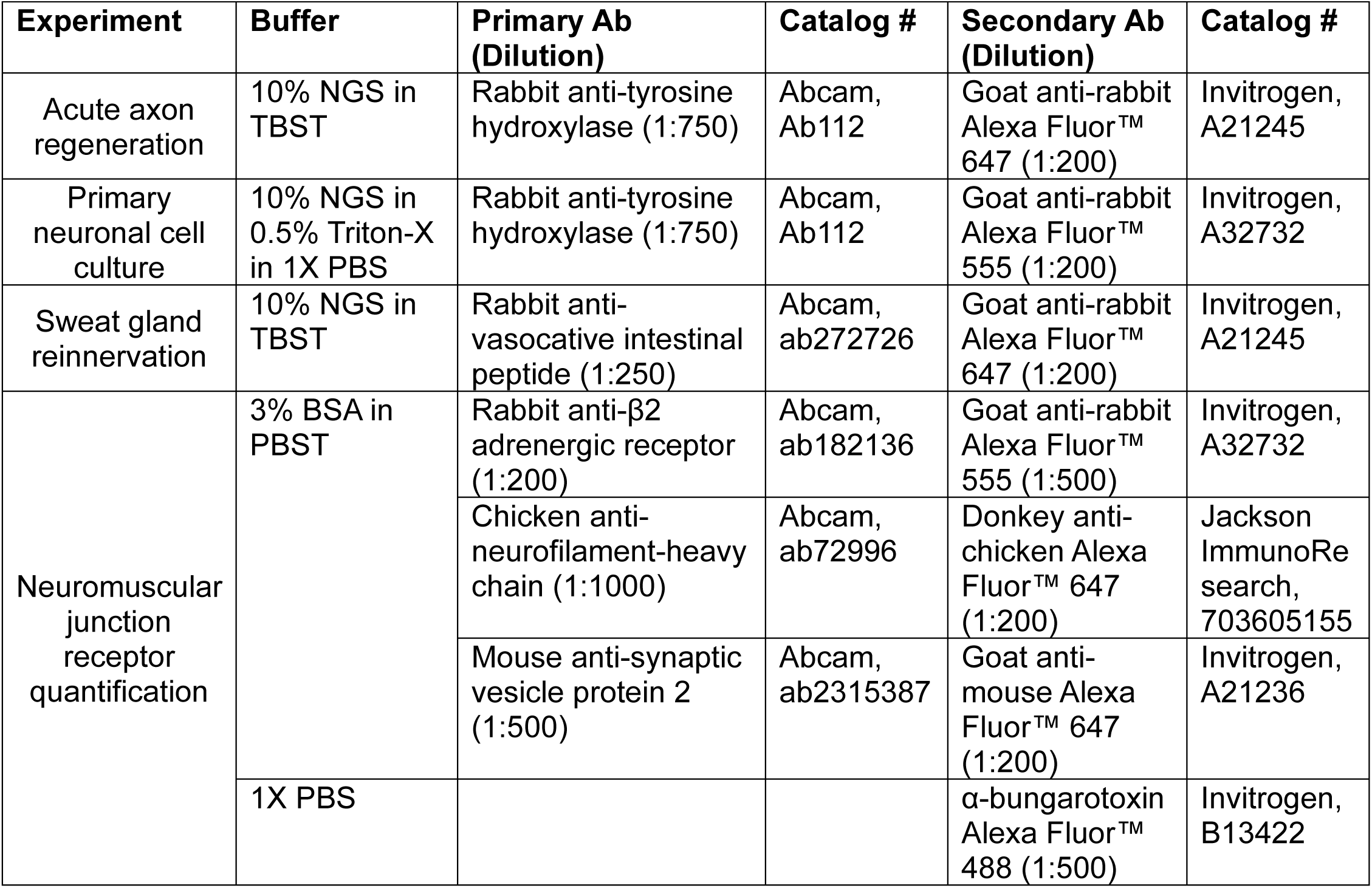
List of antibodies. Comprehensive list of antibodies used in each experiment with their dilutions, catalog numbers, and buffers. NGS = normal goat serum. TBST = 1X Tris-buffered saline with 0.1% Tween® 20 detergent. BSA = bovine serum albumin. PBST = 1X phosphate buffered saline with 0.1% Tween® 20 detergent. PBS = phosphate buffered saline.

| Experiment | Buffer | Primary Ab (Dilution) | Catalog # | Secondary Ab (Dilution) | Catalog # |
| --- | --- | --- | --- | --- | --- |
| Acute axon regeneration | 10% NGS in TBST | Rabbit anti-tyrosine hydroxylase (1:750) | Abcam, Ab112 | Goat anti-rabbit Alexa Fluor™ 647 (1:200) | Invitrogen, A21245 |
| Primary neuronal cell culture | 10% NGS in 0.5% Triton-X in 1X PBS | Rabbit anti-tyrosine hydroxylase (1:750) | Abcam, Ab112 | Goat anti-rabbit Alexa Fluor™ 555 (1:200) | Invitrogen, A32732 |
| Sweat gland reinnervation | 10% NGS in TBST | Rabbit anti-vasocative intestinal peptide (1:250) | Abcam, ab272726 | Goat anti-rabbit Alexa Fluor™ 647 (1:200) | Invitrogen, A21245 |
| Neuromuscular junction receptor quantification | 3% BSA in PBST | Rabbit anti-β2 adrenergic receptor (1:200) | Abcam, ab182136 | Goat anti-rabbit Alexa Fluor™ 555 (1:500) | Invitrogen, A32732 |
|  |  | Chicken anti-neurofilament-heavy chain (1:1000) | Abcam, ab72996 | Donkey anti-chicken Alexa Fluor™ 647 (1:200) | Jackson ImmunoResearch, 703605155 |
|  |  | Mouse anti-synaptic vesicle protein 2 (1:500) | Abcam, ab2315387 | Goat anti-mouse Alexa Fluor™ 647 (1:200) | Invitrogen, A21236 |
|  | 1X PBS |  |  | α-bungarotoxin Alexa Fluor™ 488 (1:500) | Invitrogen, B13422 |

### Surgical procedures

Prior to the start of surgery, animals received 5 mg/kg of 1.5 mg/ml oral suspension meloxidyl (Pivetal, ANADA 200-550) for postoperative analgesia. Mice were induced under anesthesia with 3% isoflurane in 1 L/min oxygen and maintained at 2% isoflurane in 1 L/min oxygen. Using a small (about 1 cm) skin incision along the femur, the sciatic nerve was exposed by blunt dissection between the hamstring muscles.

The acute No Treatment group received a transection of the sciatic nerve at the level of the midthigh with sharp, angled microscissors (Fine Science Tools, item no. 15006-09) followed by immediate repair with fibrin glue composed of fibrinogen (Sigma-Aldrich, F387910 mg in 100 μl DI water) and thrombin (MP Biomedicals, SKU 820361, 25 μl of 1 KU/1 ml normal saline with 975 μl 45 mM CaCl2) in a 2:1 ratio, as previously described **(Figure 1A)** (de Vries et al., 2002; Thompson et al., 2014; Wariyar et al., 2022). For the acute ES group, electrical pulses were delivered to the nerve through 2 monopolar needle electrodes (Ambu, #74325-36/40) placed on either side of the sciatic nerve at 20 Hz, 0.1 ms pulse duration, for 1 hour at a supramaximal stimulus intensity at the time of the nerve transection and repair **(Figure 1B)** (Brushart et al., 2002; English et al., 2007; Thompson et al., 2014; Park et al., 2019). For the acute CL group, a conditioning crush lesion was made at the level of the midthigh in the form of 3 consecutive 30-second crushes with a pair of fine forceps (Navarro and Kennedy, 1990a). One week later, the sciatic nerve was transected (test lesion) at the site of the conditioning lesion followed by immediate repair with fibrin glue **(Figure 1C)**. The hamstring muscles were closed with 5-0 absorbable suture (CP Medical, 421A) prior to skin closure with 5-0 nylon suture (MYCO Medical Supplies, N661-SI). All operations were unilateral because the conditioning lesion has been reported to promote regeneration of the contralateral sciatic nerve (Yamaguchi et al., 1999; Ryoke et al., 2000). A separate cohort of WT mice received unilateral transection and repair of the sciatic nerve and randomized into No Treatment, CL, or ES groups for acute fluorescent retrograde tracing experiments **(Figure 1D)**.

For the BL-OG *in vivo* studies, ThCre (control) and ThCre:eLMO3 mice received bilateral transections of the sciatic nerve followed by repairs with fibrin glue. Immediately after the repairs, CTZ (NanoLight Technology, #303-INJ) was administered intraperitoneally (IP) at 10 mg/kg body weight with a 25-gauge 1 ml syringe (BD, 309626), consistent with previous studies **(Figure 1E)** (Crespo et al., 2021; English et al., 2021; Ecanow et al., 2022).

To study long-term regeneration, the sciatic nerve was exposed and electrically stimulated as described above. In the Sham group, the sciatic nerve was exposed, and the electrodes were placed for 1 hour, but no stimulation was delivered. To prevent sprouting of an intact nerve into the denervated footpads, the ipsilateral saphenous nerve was ligated with 8-0 nylon suture, and a distal segment was removed to prevent regeneration **(Supplemental Figure 2)** (Kinnman, 1986; Navarro and Kennedy, 1988, 1989; Ahčan et al., 1998). All animals in the long-term regeneration assays were anesthetized for equal amounts of time and received a unilateral injury.

After surgery, all mice were placed in a clean cage on a warming pad and observed every 15 minutes until they became ambulatory. They received 5 mg/kg of oral meloxidyl every day for 3 days after surgery.

### Quantification of acute axon regeneration

Two weeks after transection and repair of the sciatic nerve in the No Treatment, CL, ES (all YFP-16 mice), ThCre, and ThCre:eLMO3 groups, the sciatic nerves were collected from the mice while the animals were under anesthesia. The nerves were postfixed for 30 minutes with 4% paraformaldehyde (PFA) (Sigma-Aldrich, P6148) in 1X PBS (0.01M phosphate buffered saline, Sigma-Aldrich, P4417) at room temperature (RT) prior to being placed in 20% sucrose with 0.02% sodium azide in 0.1M PBS and allowed to sink at 4°C. The nerves were then cryosectioned longitudinally at 20 µm and placed on charged slides (VWR Superfrost® Plus, 48311-703). The YFP-16 mice in the No Treatment, CL, and ES groups were euthanized after sciatic nerve collection via IP injection of 150 mg/kg 10 mg/ml Euthasol® (Virbac AH Inc, ANADA 200-071, NDC 051311-050-01) in 0.9% bacteriostatic saline (Hospira, Inc., NDC 0409-1966-02).

The longitudinal sciatic nerve sections were blocked with 10% normal goat serum (NGS) (VWR International, GTX73206) in 1X Tris-buffered saline (Pierce, 28358) with 0.1% Tween® 20 detergent (TBST) (Fisher Scientific, BP337500). Because tyrosine hydroxylase (TH) is the rate-limiting enzyme for norepinephrine the major neurotransmitter for postganglionic sympathetic neurons (Molinoff and Axelrod, 1971), synthesis, a rabbit anti-TH antibody was used to visualize sympathetic axons. Slides were incubated at RT overnight in rabbit anti-TH (Abcam, ab112) diluted 1:750 in the blocking buffer, washed 3x 10-minute with TBST, incubated in goat anti-rabbit Alexa Fluor™ 647 (1:200, Invitrogen, A21245) in the blocking buffer at RT for 2 hours, and washed 4x 10-minute with TBST. After drying, the slides were mounted with Fluoro-Gel with Tris (Electron Microscopy Sciences, 17985-10) and were allowed to completely harden prior to being imaged on a Nikon Ti-E fluorescent microscope on the 10x objective.

In Fiji, linear regions of interest (ROIs) were drawn that spanned the width of the nerve section every 250 μm starting from 250 μm proximal to the injury site, identified by the disorderly growth of YFP+ axons. The Cell Counter plugin was used to manually count the number of YFP+ and TH+ axons that crossed each linear ROI. The number of axons that crossed at each ROI was divided by the number of axons that crossed the ROI 250 μm proximal to the injury site to calculate a regenerative index. For each animal, 2-5 longitudinal nerve sections were analyzed that were at least 40 μm apart.

The regenerative indices of the technical replicates were averaged for each biological replicate.

### Bioluminescence quantification

The bilateral lumbar sympathetic ganglia were removed from 4 eLMO3 and 4 ThCre:eLMO3 mice that were exsanguinated with 1X PBS. The ganglia were immediately placed in ACSF (84 mM NaCl, 2.5 mM KCl, 1.2 mM NaH2PO4, 26 mM sucrose, 75 mM glucose, 4 mM MgSO4, 0.5 mM CaCl2) in a 96-well plate. The 96-well plate was placed in luminescence plate reader, and broad-spectrum luminescence was recorded continuously. Four wells with only ACSF and no tissue were used as a “blank” control. CTZ was added to all wells at a 50 μM concentration at timepoint 0 (English et al., 2021). Luminescence recordings were continued for 3 hours. Area under the curve analysis was performed on all wells.

### Local field potential recordings

The bilateral sympathetic ganglia were dissected from exsanguinated eLMO3 and ThCre:eLMO3 mice and pinned tautly in a dish with 1 mg/ml dispase/collagenase with 2.5 mg/ml papain in DMEM for 1 hour at 37°C. After digestion, the enzymes were washed off and replaced with ACSF. The recording electrode was placed within the ganglia, and blue light (480 nm wavelength) pulses were administered. Keeping the electrode in place, the ganglia were bathed in 50 μM CTZ in ACSF after the light pulses.(English et al., 2021) The change in the extracellular field potentials in response to these interventions was quantified.

### Lumbar sympathetic neuronal cell culture

To prepare for primary neuronal cell culture, 100 μl of laminin (Gibco, 23017015) was combined with 1.1 ml of a poly-D-lysine (PDL) in Hank’s balanced salt solution (HBSS) mixture (100 μl of PDL (Sigma, P6407) in 10 ml of HBSS (Corning, 21-023- CM)). Flame-sterilized glass coverslips (12 mm) were coated with the 100 μl of the laminin-PDL solution at least 1 hour prior to cell seeding.

For each round of cell culture, the bilateral lumbar sympathetic ganglia from 1 eLMO3 (control) and 1 ThCre:eLMO3 cage mate were extracted on the same day. The mice were deeply anesthetized under 5% isoflurane in 1 L/min oxygen. When no response was elicited by noxious stimuli, cervical dislocation was performed prior to rapid exsanguination with 1X PBS. Both mice were then transferred to a cooled dissection tray, and dissection was performed within a semi-enclosed ventilated hood dissection table.

Ganglia were placed in 1 ml RT HBSS prior to being digested in 1 ml dispase/collagenase (EMD Millipore, SCR140, resuspended at 1 mg/ml with 1% fetal bovine serum in NB-A) at 37°C for 1 hour. After 1 hour, the dispase/collagenase was removed and 100 μl of DNase (Worthington, LK003172) was added to the ganglia for 2.5 minutes at 37°C. Then, 800 μl of 37°C HBSS was added to the ganglia without removing the DNase. The ganglia were dissociated using progressively narrower plastic pipette tips prior to being spun down at 3000 rpm for 3 minutes.

The supernatant was removed, and 50 μl of 37°C Neurobasal-A (NB-A) Media (Gibco, 10888-22) was used to resuspend each pellet. The laminin-PDL coating was aspirated off the glass coverslips, and the resuspended cells were seeded to the coverslips followed by 450 μl of Neurobasal-A Complete (96% NB-A, 1% GlutaMAX (Gibco, 10888-022), 1% penicillin-streptomycin (Sigma, P4333), 2% 50x B27 (Gibco, 17504044)) with 50 μM CTZ (English et al., 2021). The cells were allowed to incubate for 1 hour at 37°C with CTZ in the media prior to 3 washes with NBA Complete. After the 3 washes, the cells were then incubated for another 48-72 hours at 37°C before the media was aspirated and replaced with 4% PFA. The cells were postfixed for 15 minutes followed by 3 10-minute washes with 1X PBS.

For immunofluorescence, cells were permeabilized with 0.5% Triton-X in 1X PBS for 15 minutes followed by 1 hour blocking with 10% NGS in 0.5% Triton-X in 1X PBS for 1 hour at RT. Rabbit anti-TH (1:750) was diluted in the blocking buffer and allowed to incubate on the cells overnight. The cells were washed 3 times for 10 minutes with PBS prior to applying goat anti-rabbit Alexa Fluor™ 555 (1:200, Invitrogen, A32732) for 2 hours. Following 3 more 10-minute washes with 1X PBS, the coverslips were mounted on glass slides with Fluoro-Gel with Tris. All slides were imaged on a Nikon Ti-E fluorescent microscope on the 20x objective.

All neurons with a visible cell body and nuclear shadow with an enclosing radius of at least 100 μm were imaged and traced using the SNT toolbox within the Neuroanatomy plugin in Fiji (Arshadi et al., 2021). Sholl analysis was performed, and the enclosing radius and branching index of each neuron were determined **(Figure 1F)**. In order for an animal to be counted for the final analysis, at least 8 traceable neurons needed to be visible. Values from neurons derived from the same mouse were averaged together. Animals dissected on the same day were paired. A total of 65 eLMO3 neurons and 43 ThCre:eLMO3 neurons were traced and analyzed.

### Fluorescent retrograde tracing

To further evaluate axon regeneration in the acute regeneration groups, nerve soak in fluorescent retrograde tracer was performed. In the WT mice that were randomized to either No Treatment, CL, or ES groups and the ThCre and ThCre:eLMO3 mice, the injured sciatic nerves were re-exposed 2 weeks after injury under anesthesia and transected 5 mm distal to the original injury site. The proximal stump was placed in fast blue (FB) retrograde tracer (1 mg/ml deionized water, Polysciences, 17740) for 1 hour. Vacuum silicone grease (Oakwood Chemical, 106690) was used to create a barrier around the soak site to prevent tracer contact with shorter regenerated axons. After 1 hour, the muscles and skin were reapproximated.

To determine the number of motor and sympathetic neurons whose axons had successfully regenerated to distal muscle targets 12 weeks after unilateral sciatic nerve injury, the mice were anesthetized, and the TA and the intrinsic muscles of the foot were injected with fluorescent retrograde tracer, bilaterally, creating a contralateral “Intact*”* group. The TA was injected with 2 μl of Cholera Toxin Subunit B Alexa Fluor™ 555 (CTB555) retrograde fluorescent tracer (500 mg/1 ml deionized water, Invitrogen, C22843) distributed in at least 2 locations in the muscle belly. The intrinsic muscles of the foot were injected with 0.5 μl of FB in 3 sites (1.5 μl total), entering through the 3 web spaces between the 2^nd^ and 5^th^ toes. All injections were delivered via a 26-gauge Hamilton syringe.

The mice were euthanized 3 days after nerve soak or muscle injection, to allow for retrograde transport of the fluorescent tracer, with IP injection of 150 mg/kg Euthasol solution. Once the mice displayed no response to noxious stimuli, the animals were quickly exsanguinated with 0.9% NaCl then perfused with 4% PFA.

The L2-L5 level lumbar sympathetic ganglia, which traverse from the level of the left renal artery and the bifurcation of the aorta (Zheng et al., 2017; Tian and Ward, 2024), were collected from the mice that received a nerve soak and muscle injections. The lumbar level spinal cord was also collected from the mice that received muscle injections. The lumbar sympathetic chain was mounted directly onto glass slides with Fluoro-Gel mounting media with Tris buffer, and images were taken with a Nikon Ti-E fluorescent microscope using the 20x objective. SCs were placed in 20% sucrose in 0.1 M PBS and allowed to sink overnight at 4°C prior to cryosectioning. SCs were sectioned at 50 μm longitudinally and placed onto glass slides, and all sections were imaged using the 10x objective.

In order for a neuron to be counted as retrogradely-traced, the tracer must fill the soma and extend into the cell’s proximal neurites, and the nuclear shadow of the neuron must be visualized. If the labeled neuron did not meet these criteria, it was not counted.

### Pilocarpine sweat assay

For the long-term regeneration cohort, the recovery of the sweating response was evaluated with pilocarpine (Sigma-Alrich, P6503), a muscarinic agonist which stimulates sweating, which has been used in previous studies to track functional sympathetic regeneration as well as to characterize diabetic neuropathy (Navarro et al., 1988; Navarro and Kennedy, 1988, 1989, 1990b; Liu et al., 2017). With the animal under anesthesia, the hind paws were secured to the surgical tabletop with tape and coated with betadine using a paintbrush. Once the betadine was dry, 10% potato starch (Judee’s) in castor oil (Seven Minerals) was painted on the foot. Finally, 0.25 μl/g body weight of 1% pilocarpine in normal saline was injected subcutaneously. Eight minutes post-injection, photos of the plantar foot were taken. This assay was performed prior to the nerve injury to determine a baseline sweating response then every 2 weeks for 12 weeks. As mentioned in the “Surgical procedures” section, the ipsilateral saphenous nerve was also ligated to prevent the sprouting of intact fibers into the denervated areas of the foot **(Supplemental Figure 2)** (Kinnman, 1986; Navarro and Kennedy, 1988, 1989; Ahčan et al., 1998). Sweating droplets were identified as dark blue-black spots, due to the precipitates created in the reaction of starch to iodine, located on the footpads. This method has been shown to yield results comparable to those obtained using silicone imprints (Provitera et al., 2010).

### Sweat gland reinnervation

At 12 weeks post-injury, mice were anesthetized prior to dissection and removal of plantar foot skin bilaterally. The plantar skin tissues were postfixed in 4% PFA for 30 minutes prior to being transferred to 20% sucrose in 0.1M PBS for cryoprotection overnight at 4°C. Short axis cross sections of the plantar skin were made at 20 μm, sectioning distal to proximal, and placed on charged slides. The most distal footpad, labeled as footpad 3 **(Figure 7A)** was isolated, numbered according to previously established standards (Koike et al., 2021), and reinnervation of the sweat glands in footpad 3 was evaluated via IHC. Footpad sections were blocked with 10% NGS in TBST for 1 hour in a humidity chamber. Rabbit anti-vasoactive intestinal peptide (anti-VIP, 1:250, Abcam, ab272726) was diluted in the blocking buffer and allowed to incubate on the footpad sections overnight at RT. Following 3 10-minute washes with TBST, goat anti-rabbit Alexa Fluor™ 647 (1:200) in the blocking buffer was allowed to incubate at RT for 2 hours followed by 4x 10-minute washes with TBST. VIP is a neuropeptide that is present in cholinergic sympathetic and parasympathetic fibers (Lundberg et al., 1980; Lundberg et al., 1981b; Lundberg et al., 1981a; Lundberg et al., 1985; Lundberg et al., 1987; Lundberg et al., 1991), and it modulates the cholinergic response of sweat glands while having vasodilator properties (Yamashita et al., 1987). Furthermore, functional sweat output is associated with VIP immunoreactivity (Levy et al., 1992; Navarro et al., 1997). Therefore, VIP+ axons were used to quantify sympathetic reinnervation. Because YFP expression in postganglionic sympathetic axons is negligible in YFP-16 mice **(Supplemental Figure 1)** (Feng et al., 2000), YFP+ axons were used to quantify aberrant reinnervation.

VIP+ and YFP+ axons were manually traced in Fiji in the sweat gland area. The total length of the axons was then divided by the sweat gland area. Three footpad sections at least 20 μm apart were analyzed and averaged.

### High performance liquid chromatography for purine quantification

In the long-term regeneration group with unilateral injury, the bilateral gastrocnemii muscles were collected while the mice were under anesthesia, therefore creating an “intact” group. The muscle was split into 4 approximately equal parts and weighed prior to flash freezing with liquid nitrogen. A quarter of the lateral head of the GA was then powdered using a mortar and pestle, nestled within dry ice, adding liquid nitrogen to the mortar as necessary. Once powdered, the muscle was transferred to cold 0.2 M perchloric acid (Sigma-Alrich, 311421) at a concentration of 1 ml/10 mg of muscle(Weicker et al., 1990), and the solution was vortexed and flash frozen in liquid nitrogen.

After thawing on ice, the samples were spun down for 5 minutes at 3000 rpm. The pellet was resuspended in 1 ml 2% sodium dodecyl sulfate (SDS) and sonicated for approximately 1.5 minutes while ensuring that the solution was not warmed. Protein content was quantified using the Pierce BCA assay (Thermo Scientific, 23227).

From the supernatant of each sample, 250 μl was removed and placed in a new tube, to which 3.5% v/v 3.5 M K2CO3 (Oakwood Chemical, 102941) was added. The samples then sat on ice for 30 minutes prior to being centrifuged at 13000 rpm for 15 minutes at 4°C. The supernatants were then transferred to 0.2 μm centrifuge filters (Thermo Scientific, F2517-5) and spun at 8000 rpm for 5 minutes at 4°C. The filtrate was then transferred to high performance liquid chromatography (HPLC) vials for Waters HPLC autosampler (Waters Quality Parts, WAT022476), and purines were quantified by Emory University’s HPLC core facility. Purine quantifications were then normalized to protein content (Sabina et al., 1983). The cellular energy charge was calculated using the following equation:

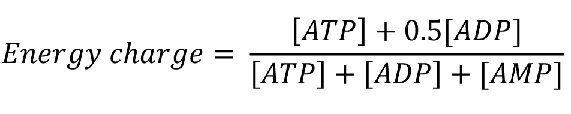

### Electromyography and muscle collection

Fine wire electromyography (EMG) was performed bilaterally, as previously described (Wariyar et al., 2022), on the gastrocnemius 12 weeks after unilateral sciatic nerve transection and repair in a separate cohort of YFP-16 mice. Briefly, fine wire electrodes (A-M Systems, 762000) were placed into the gastrocnemius muscle to record the compound muscle action potentials (CMAPs) evoked by 0.1 ms electrical pulses with incrementally increasing stimulus intensities with the animal under anesthesia. The ramp was discontinued once the rectified CMAP amplitude plateaued. The maximum functional motor reinnervation of the muscle is presumed to be correlated with the maximum CMAP amplitude.

After completing the EMGs, the bilateral triceps surae muscle groups, composed of the medial and lateral heads of the gastrocnemius, soleus, and plantaris (Dalmau- Pastor et al., 2014), were dissected from the animal, tendon-to-tendon, bilaterally.

Excess Achilles tendon was removed before weighing the triceps surae. The mass of the injured side was compared to the mass of the intact side in each animal.

### Neuromuscular junction receptor quantification

Twelve weeks after unilateral sciatic nerve transection and repair, the gastrocnemius muscles were collected bilaterally, quartered, and flash frozen in liquid nitrogen as detailed above. A quarter of the lateral head of the gastrocnemius from each leg of each mouse was then sectioned longitudinally at 30 μm and placed on charged slides. Muscle sections were then postfixed with 4% PFA in 1X PBS for 5 minutes at RT followed by 3 5-minute washes with 1X PBS. In a humidity chamber, the sections were then blocked with 3% bovine serum albumin (BSA, Sigma-Aldrich, A2153) in 1X PBS with 0.1% Tween® 20 detergent (PBST) for 1 hour at RT followed by the primary antibodies rabbit anti-β2 adrenergic receptor (ADRB2, 1:200, Abcam, ab182136), chicken anti-neurofilament-heavy chain (NF-H, 1:1000, Abcam, ab72996), and mouse anti-synaptic vesicle protein 2 (SV2, 1:500, Abcam, ab2315387) in blocking buffer overnight at RT. NF-H and SV2 were used to visualize the motor reinnervation because the YFP had been quenched in the flash frozen muscles. The slides were then washed 3x with PBST, 10 minutes each wash, then the following secondary antibodies were added to the slides in the blocking buffer: goat anti-rabbit Alexa Fluor™ 555 (1:500), donkey anti-chicken Alexa Fluor™ 647 (1:200, Jackson ImmunoResearch, 703605155), and goat anti-mouse Alexa Fluor™ 647 (1:200, Invitrogen, A21236). The secondary antibodies were allowed to incubate at RT for 2 hours prior to 3x 10-minute washes with PBST. Alpha-bungarotoxin Alexa Fluor™ 488 (BTX, 1:500, Invitrogen, B13422) in 1X PBS was added to the slides for 30 minutes and washed off with 3x 10-minute 1X PBS washes.

NMJs were identified with the BTX channel (FITC) and were imaged using the 60x oil objective on a Nikon Ti-E fluorescent microscope. In Fiji, max projections of individual NMJs were created, and for each NMJ, a region of interest (ROI) was identified in the BTX channel using the Threshold function in Fiji. The ROI was further refined using the Selection Brush Tool to only encompass the BTX+ area. Three rectangular ROIs approximately the size of the NMJ were drawn randomly in the BTX- negative areas to determine background fluorescence. The average value of the 3 background ROIs was found. The average fluorescence intensity was found for both the BTX (representative of acetylcholine receptors) and ADRB2 (representative of β2- adrenergic receptors) channels at the NMJ as well as in the 3 background ROIs. Finally, the fluorescence values of the NMJ ROI for BTX and ADRB2 were divided by the average background of their respective channels. The area of the NMJ was also recorded. At least 40 NMJs were analyzed per mouse: 251 Intact (n = 5), 204 Sham (n= 4), and 199 ES (n = 4). A total of 654 NMJs were incorporated into this assay.

### Bulk RNA sequencing of the lumbar sympathetic ganglia

Male WT mice from were used for bulk RNA sequencing of the lumbar sympathetic ganglia. All mice were between 8-9 weeks of age at the time of the start of the experiment and weighed between 20-27 g. The animals were randomized into Intact (1 hr with nerve exposure and no stimulation), Sham (1 hr of sham stimulation), and ES groups, with Sham and ES groups receiving bilateral sciatic nerve transections.

Therefore, all animals received the same amount of anesthesia (3% isoflurane in 1 L/min oxygen for induction, 2% isoflurane for maintenance) and the same amount of preoperative and postoperative analgesia (5 mg/kg meloxicam prior to surgery and once a day for 3 days after surgery). The incisions were closed as described above.

Three days after sciatic nerve transection(Shin et al., 2019; Kalinski et al., 2020; Senger et al., 2021), the mice were deeply anesthetized with 5% isoflurane in 1 L/min oxygen, and upon no response to noxious stimuli, such as forepaw pinch, cervical dislocation was performed. The animals were then rapidly exsanguinated with 1X PBS followed by 15 seconds of 30% RNA*later* (Invitrogen, AM7020) in 1X PBS at a flow rate of 8 ml/min. The bilateral lumbar sympathetic ganglia were then collected and placed in 100% RNA*later™*, combining the ganglia from 2 mice for one sample to achieve a sufficient quantity of RNA. Each group required 8 mice for an n of 4 per group.

The total RNA was extracted from the pooled ganglia for each sample within 2 days of ganglia collection. The tissues were transferred to RNase-free pink bead lysis kits (Midsci, PINKE1-RNA) and lysed using a Bullet Blender® set at 12 speed for 30 seconds. The samples were processed using RNAeasy® Micro Kits (Qiagen, 74004) as specified by manufacturer instructions, but an extra wash with buffer RPE (Qiagen, 1018013) was added to all samples to optimize the 260/230 ratio. The total RNA was eluted in 14 μl of RNase-free water (Research Products International, 248700).

RNA quantity and purity were assessed using a NanoDrop™ One Microvolume UV-Vis Spectrophotometer, and all samples had RNA concentrations of at least 30 ng/μl and 260/280 ratios between 2.0-2.1. The samples were then flash frozen in liquid nitrogen prior to being sent for bulk mRNA-sequencing by Novogene Co. (CA, USA), which performed the library construction, quality control, sequencing, and identification of differentially expressed genes (DEGs). A DEG was considered significant if the - log10(p-value) was greater than or equal to 2 and was considered upregulated if the log2(fold change) was greater than or equal to 1.5 and downregulated if the log2(fold change) was less than or equal to 1.5 **(data not shown, see Data Availability)**.

### Statistical analysis

Sample sizes for each experiment were calculated to reach 80% power and β < 0.2 based on previously published work and preliminary experiments (Liu et al., 2017; Wariyar et al., 2022). Two-way RM ANOVAs were performed for assays that evaluated 2 or more groups over distances or over time (i.e. regenerative indices and sweating response). For the *in vitro* experiments, which involved the neurons from cage mates being cultured on the same day, two-tailed paired t-tests were performed. For assays that involved more than 2 groups, ordinary one-way ANOVAs were performed followed by post hoc Tukey’s multiple comparisons test. If SDs between groups were determined to be significantly different from each other by the Brown-Forsythe test, Brown-Forsythe ANOVA tests were performed followed by post hoc Tamhane T2 multiple comparisons test or Games-Howell’s multiple comparisons test if n > 50. If the outcomes were nonparametric (i.e. discrete cell counts) or if the residuals failed normality testing via the Anderson-Darling test, the Kruskal-Wallis test was performed followed by post hoc Dunn’s multiple comparisons test. If the nonparametric data only contained 2 groups, the Mann-Whitney test was performed. To determine if the number of retrogradely labeled neurons and GA CMAPs had returned to baseline levels, the Intact group was created from the contralateral uninjured side for Figure 9, 10, and 12, so the experimental groups could be expressed as a percentage of Intact. For statistical analysis to be performed on the Intact group, the means of the raw neuron counts and CMAP amplitudes were found, and each raw value was divided by their respective means to accurately reflect the variability of these assays while keeping the overall mean of the Intact group at 100%.

Area under the curve (AUC) analysis was performed for the 3 hours of recording bioluminescence from the lumbar sympathetic ganglia of BL-OG mice after introducing CTZ. Ordinary one-way ANOVA followed by post hoc Tukey’s multiple comparisons test was performed to evaluate for significance. LFP recordings were unpaired, so two-tailed unpaired t-tests were used for statistical analysis.

HPLC results were analyzed via estimation statistics using a shared control (Intact) (Ho et al., 2019) Power analysis had determined that the group sizes needed to be increased from n = 7 to n = 19 to achieve statistical significance; therefore, to reduce the number of animals needed in this study, we opted to use estimation statistics, which will allow for more clear interpretation of the effect sizes. All statistical analyses were performed using Graphpad Prism or www.estimationstats.com /#/ (Ho et al., 2019).

Comparisons were determined to be significant if p < 0.05. GraphPad Prism, Microsoft PowerPoint, and BioRender.com were used to generate the figures.

## RESULTS

*Neither conditioning lesion nor electrical stimulation enhances acute sympathetic axon regeneration* Peripheral nerves regenerate slowly, and the ability of axons to regenerate and the supportive function of Schwann cells decline over time without additional interventions such as ES or CL (Gutmann et al., 1942; Sunderland, 1947). To test the effect of perioperative ES on sympathetic axon regeneration, regenerative indices were calculated for motor and sensory axons (represented by YFP in the YFP-16 mice) and sympathetic axons (represented by TH staining). A CL paradigm, shown to enhance motor and sensory regeneration independent of the axonal electrical properties, was used as a comparison group.

Two weeks after transection and repair of the sciatic nerve, both the CL and ES enhanced motor and sensory regeneration (2-way RM ANOVA, experimental group factor F (2, 19) = 3.872, p = 0.0388, post hoc Tukey’s multiple comparisons test) starting at 1750 μm distal to the injury site (1750 μm ES vs No Treatment p = 0.0446; 2000 μm CL vs No Treatment p = 0.0131, ES vs No Treatment p = 0.0472; 2250 μm CL vs No Treatment p = 0.0056, ES vs No Treatment p = 0.0310; 2500 μm CL vs No Treatment p = 0.0269, ES vs No Treatment p = 0.0307; 2750 μm CL vs No Treatment p = 0.0199; 3000 μm p = 0.0049) **(Figure 2A-D)**. In contrast, neither intervention enhanced sympathetic regeneration (2-way RM ANOVA, experimental group factor F (2, 19) = 1.218, p = 0.3179) **(Figure 2E-H)**. Retrograde tracing via nerve soak 5 mm distal to the injury site also revealed no difference in the number of labeled sympathetic neurons between No Treatment, ES, and CL (Kruskal-Wallis test, p = 0.6955) **(Figure 2I-L)**.

**Figure 2:**
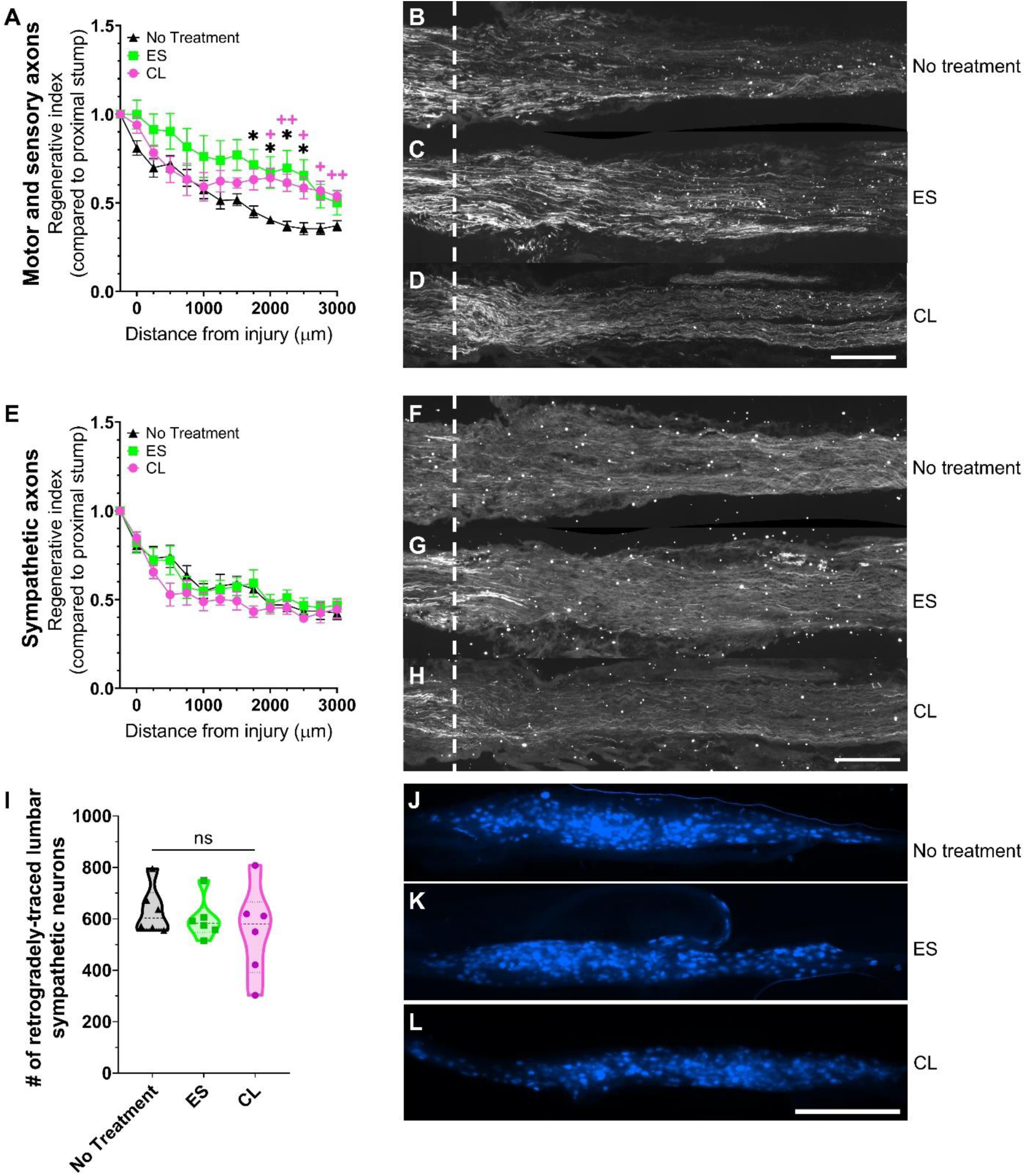
Electrical stimulation and conditioning lesion do not acutely enhance sympathetic regeneration but do enhance motor and sensory regeneration. (A) Regenerative indices of motor and sensory axons 2 weeks after sciatic nerve transection and repair in response to No Treatment (black triangles), electrical stimulation (ES, green squares), and conditioning lesion (CL, magenta circles) plotted as distance from the injury site. Data shown as mean ± SEM. Two-way RM ANOVA with post hoc Tukey’s multiple comparisons test. * = No Treatment vs. ES. + = No Treatment vs. CL. * or + p < 0.05. ++ p < 0.01. No Treatment n = 6, ES n = 8, CL n = 8. (B-D) Representative sciatic nerve sections showing motor and sensory axons visualized with yellow fluorescent protein. (E) Regenerative indices of sympathetic axons 2 weeks after sciatic nerve transection and repair in response to No Treatment (black triangles), electrical stimulation (ES, green squares), and conditioning lesion (CL, magenta circles) plotted as distance from the injury site. (F-H) Representative sciatic nerve sections showing sympathetic axons labeled by tyrosine hydroxylase immunohistochemistry (Alexa Fluor® 647). Data shown as mean ± SEM. (I) Violin plot of retrogradely-labeled lumbar sympathetic neurons that had extended their axons at least 5 mm distal to the injury site 2 weeks after injury. Data shown as median ± interquartile range. (J-L) Representative L2 sympathetic ganglia with retrogradely-traced neurons via fast blue nerve soak. Scale bars = 500 μm. n = 6 for No Treatment, ES, and CL.

### Bioluminescent activation of sympathetic neurons inhibits their growth in vivo and in vitro

The ThCre:eLMO3 mouse expresses eLMO3, a luminopsin channel that is conjugated to a luciferase enzyme **(Figure 3A)**, in TH+ neurons, which encompasses the vast majority of postganglionic sympathetic neurons. The eLMO3 channel also possesses a YFP tag, which can be visualized at the cell body as well as axons present in the lumbar sympathetic chain ganglia **(Figure 3B)**. *Ex vivo* lumbar sympathetic chain ganglia in artificial cerebrospinal fluid (ACSF) exhibited a significant long-lasting increase in bioluminescence in the presence of CTZ that was (ordinary one-wave ANOVA, F = 16.88, p = 0.0009, post hoc Tukey’s multiple comparisons test: Blank vs ThCre:eLMO3 p = 0.0018, eLMO3 vs ThCre:eLMO3 p = 0.0019) **(Figure 3C-D)**.

**Figure 3:**
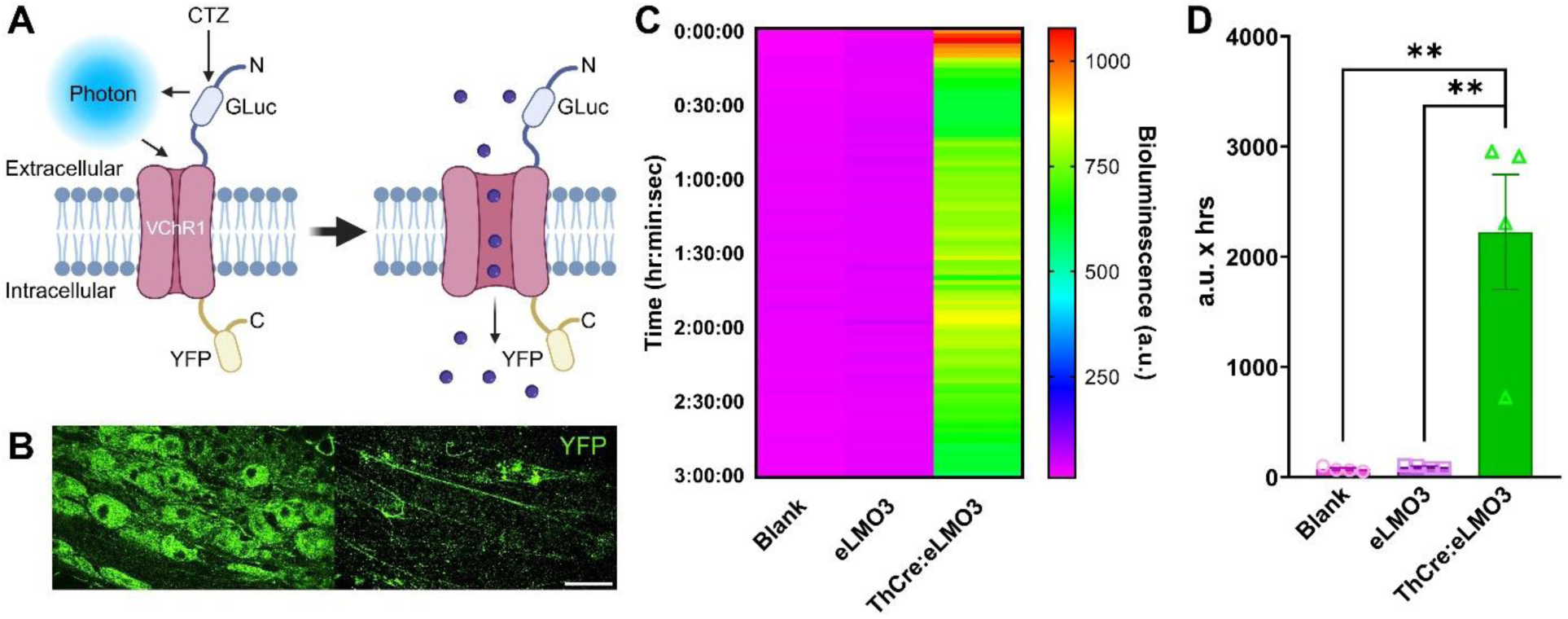
The lumbar sympathetic ganglia of ThCre:eLMO3 mice express the excitatory luminopsin eLMO3 and exhibit long-lasting bioluminescence in the presence of coelenterazine. (A) Diagram of enhanced luminopsin 3 (eLMO3), which is comprised of a luciferase enzyme from *Gaussia princeps* (GLuc) fused to a red-shifted channelrhodopsin derived from *Volvox carteri* (VChR1). The cytosolic tail is coupled to yellow fluorescent protein (YFP). The channel can be activated by the biological light that is produced when the luciferase catalyzes the substrate coelenterazine (CTZ). Adapted from Berglund et al. (2016) and English et al. (2021). (B) Presence of intracellular YFP tag in the cell bodies of lumbar sympathetic neurons and their axons visualized in the ganglia. Scale bar = 50 μm. (C) Bioluminescence emitted from the L2- L5 lumbar sympathetic ganglia from eLMO3 (control) and ThCre:eLMO3 mice after being exposed to 50 μM CTZ. The blank group consisted of empty wells with the suspending media and no ganglia. (D) Area under the curve quantification of bioluminescence over time. Data shown as mean ± SEM. One-way ANOVA with post hoc Tukey’s multiple comparisons test. ** p < 0.01. n = 4 for Blank, eLMO3, and ThCre:eLMO3.

The local field potential (LFP) of the neurons in *ex vivo* lumbar sympathetic ganglia were recorded **(Figure 4A)**. The sympathetic neurons of ThCre:eLMO3 mice responded to a flash of visible blue light with a decrease in voltage indicating cation influx into the neurons **(Figure 4B)**. There was little or no deflection in the control ganglia that did not express the eLMO3 channel. **(Figure 4D)**. This difference between eLMO3 and ThCre:eLMO3 ganglia was also seen in response to CTZ **(Figure 4F)**. The negative deflection in the LFPs of ThCre:eLMO3 ganglia in response to both external light (two-tailed unpaired t-test, p = 0.0229, t = 3.038, df = 6) and CTZ (two-tailed unpaired t-test p = 0.0290, t = 3.335, df = 4) were greater than that seen in eLMO3 (control) ganglia **(Figure 4C & E)**.

**Figure 4:**
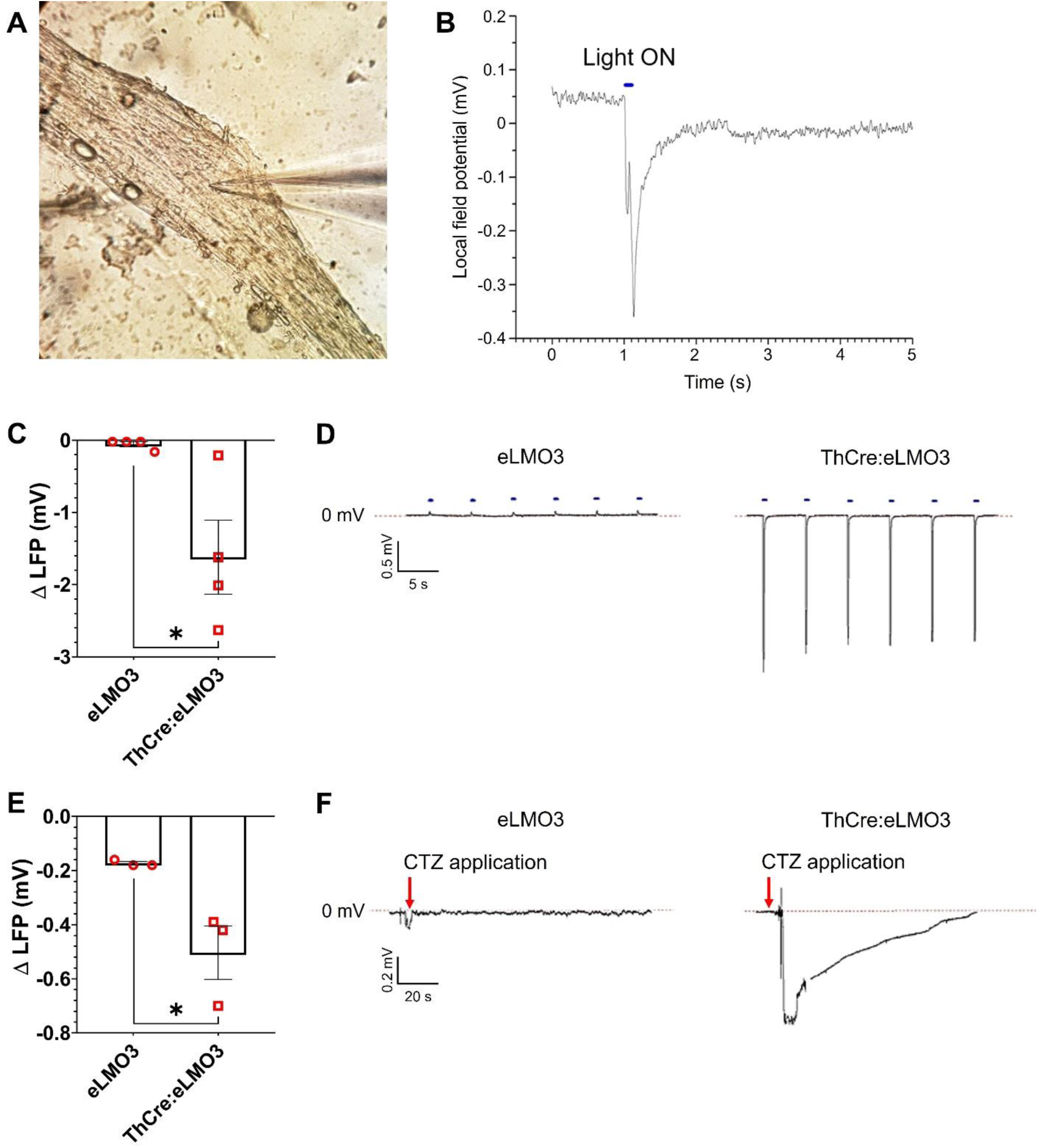
External blue light and CTZ sufficient to induce a negative deflection in the local field potential in ThCre:eLMO3 lumbar sympathetic ganglia. (A) Image of recording electrode inserted in the lumbar sympathetic ganglia. (B) Example local field potential (LFP) tracing in response to a pulse of external blue light. (C) Representative LFP tracings from eLMO3 (control) and ThCre:eLMO3 ganglia in response to pulses of external blue light. (D) Quantification of the change in LFP in response to external blue light. (E) Representative LFP tracings from eLMO3 and ThCre:eLMO3 ganglia in response to 50 μM CTZ. (F) Quantification of the change in LFP in response to CTZ. Two-tailed unpaired t-test. * p < 0.05. Data shown as mean ± SEM. eLMO3 n = 4, ThCre:eLMO3 n = 3.

Bilateral transection and repair of the sciatic nerve was performed in ThCre (control) and ThCre:eLMO3 mice with IP injection of CTZ at the time of the injury. Two weeks later, the right sciatic nerve was collected for evaluation of sympathetic regenerative indices, and the left sciatic nerve was soaked in fast blue 5 mm distal to the original injury site. There was no difference in the regenerative index between stimulated (ThCre:eLMO3) and non-stimulated (ThCre) mice (both treated with CTZ) detected by IHC up to 2500 μm distal to the injury site (2-way RM ANOVA, experimental group factor F (1, 14) = 0.03512, p = 0.8540) **(Figure 5A-C)**. However, the number of lumbar sympathetic neurons that had regenerated their axons at least 5 mm 2 weeks after injury as measured by retrograde uptake of fast blue was significantly decreased in the ThCre:eLMO3 group compared to the control group (Mann-Whitney test, p = 0.0085) **(Figure 5D-F)**.

**Figure 5:**
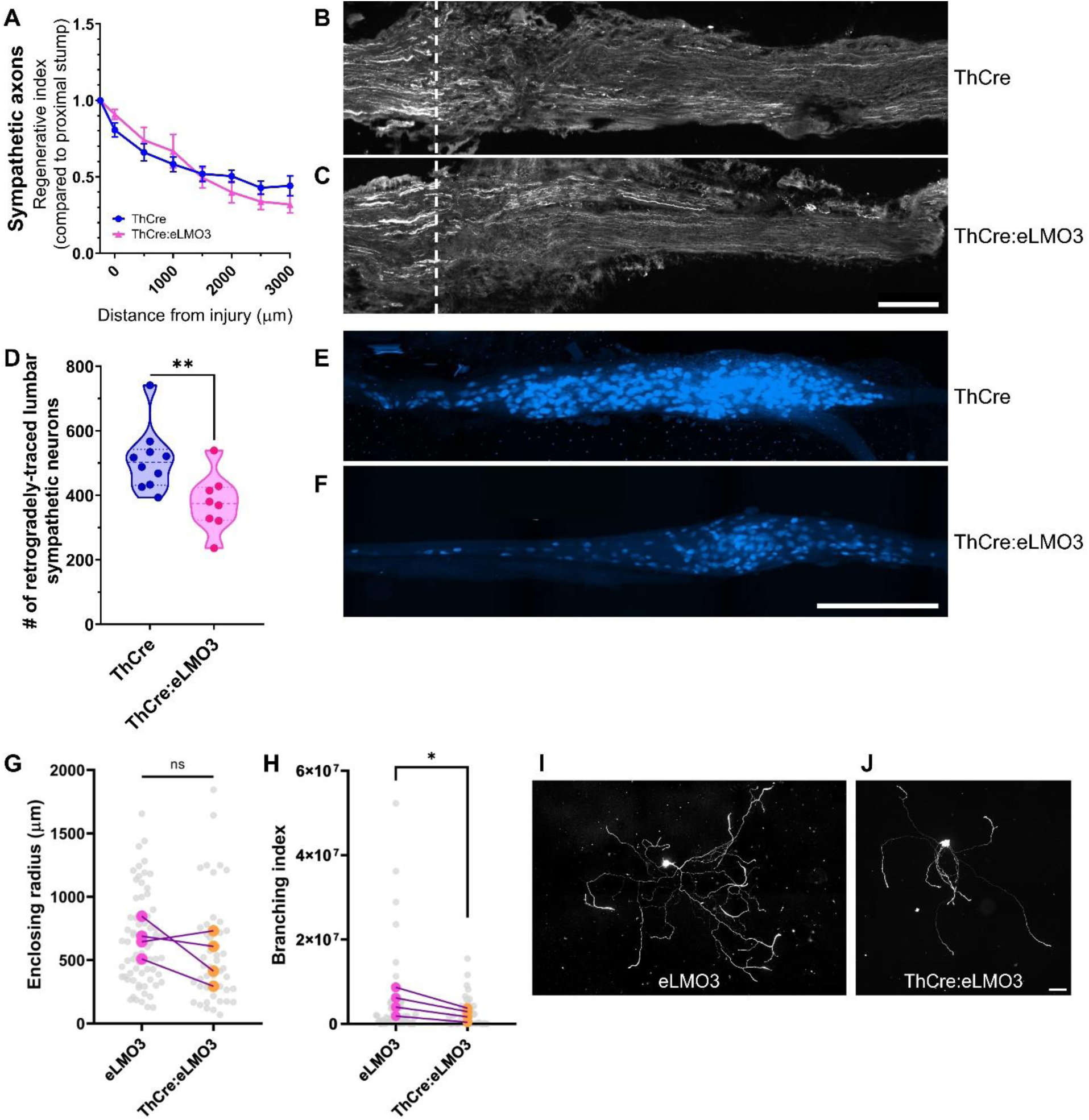
Bioluminescent optogenetic stimulation of sympathetic neurons results in impaired sympathetic regeneration *in vivo* and decreased branching of sympathetic neurons *in vitro*. (A) Regenerative indices of sympathetic axons 2 weeks after sciatic nerve transection and repair plotted as distance from the injury site. Data shown as mean ± SEM. (B-C) Representative sciatic nerve sections showing sympathetic axons labeled by tyrosine hydroxylase (TH) immunohistochemistry (AlexaFluor® 647). ThCre n = 9, ThCre:eLMO3 n = 7. (D) Violin plot of retrogradely-labeled lumbar sympathetic neurons that had extended their axons at least 5 mm distal to the injury site 2 weeks after injury. Kruskal-Wallis test with post hoc Dunn’s multiple comparisons test. ** p < 0.01. Data shown as median ± interquartile range. (E-F) Representative L2 sympathetic ganglia with retrogradely-traced neurons via fast blue nerve soak. Scale bars = 500 μm. ThCre n = 10, ThCre:eLMO3 n = 8. (G) Enclosing radii of cultured lumbar sympathetic neurons after CTZ exposure. (H) Branching indices of cultured lumbar sympathetic neurons after CTZ exposure. Two-tailed paired t-test. * p < 0.05. Representative images of lumbar sympathetic neurons in culture visualized via TH immunofluorescence (Alexa Fluor® 555). Scale bar = 100 μm. n = 4 for eLMO3 and ThCre:eLMO3.

**Figure 6:**
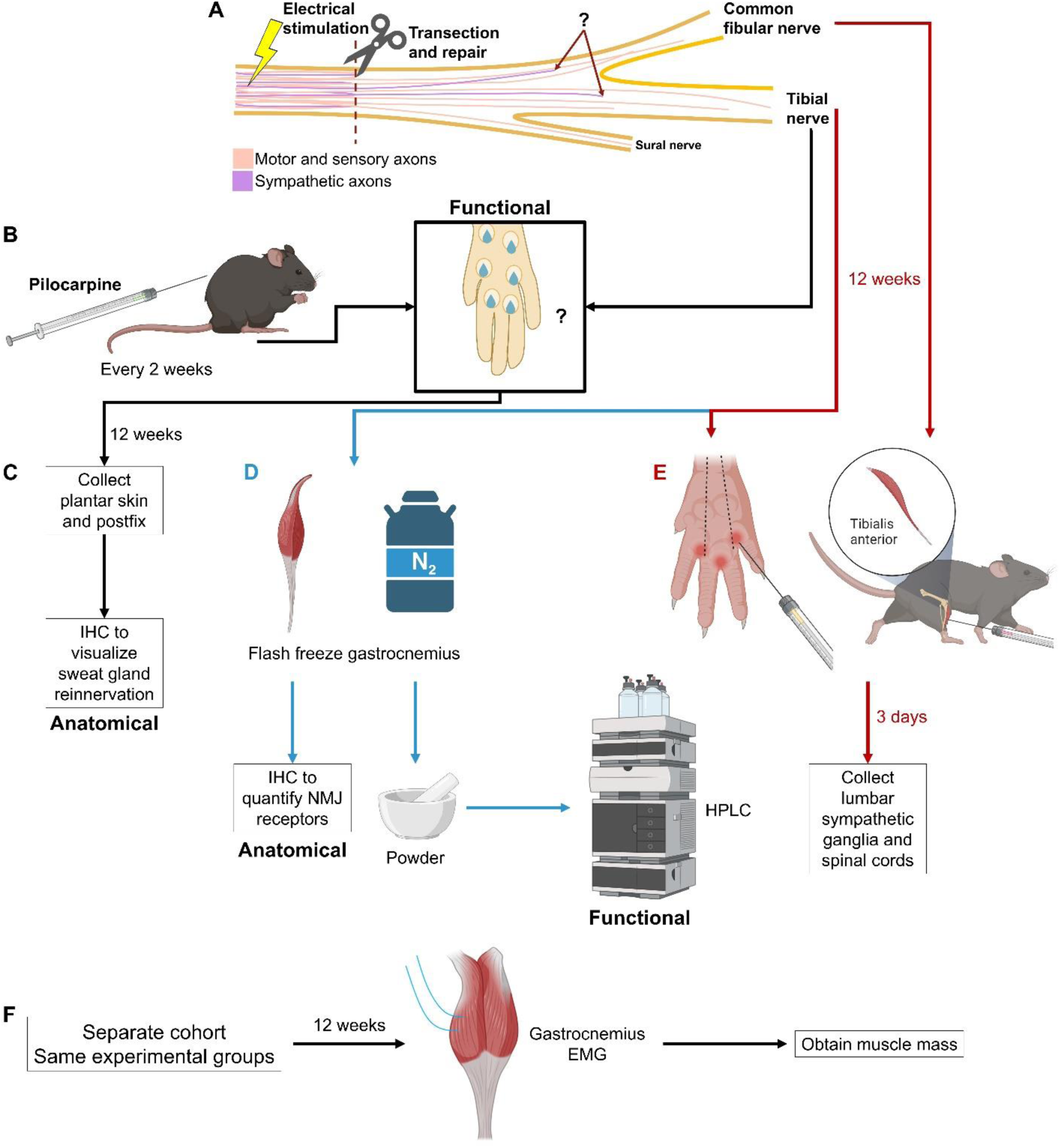
Long-term regeneration summary of methods. (A) Electrical stimulation at the time of sciatic nerve transection and repair. (B) Functional evaluation of sympathetic skin response via the pilocarpine sweat assay every 2 weeks. The plantar foot skin is innervated by the tibial nerve. (C) Evaluation of sympathetic reinnervation of the sweat glands via immunohistochemistry (IHC). (D) Evaluation of gastrocnemius neuromuscular junction (NMJ) receptor quantification via IHC and purine content after injury via high purity liquid chromatography (HPLC). (E) Injection of fluorescent retrograde tracers into the intrinsic muscles of the foot and the tibialis anterior to evaluate distal and proximal target reinnervation, reinnervation. (F) Electrophysiological evaluation via electromyography (EMG) on the gastrocnemius performed on a separate cohort of animals with measurement of muscle mass. Partially created with BioRender.com.

When eLMO3 (control) and ThCre:eLMO3 lumbar sympathetic neurons were dissociated and exposed to 1 hour of CTZ in the culture media, ThCre:eLMO3 neurons exhibited decreased branching (two-tailed paired t-test, p = 0.0284, t = 3.987, df =3) **(Figure 5H-J)**. The enclosing radius of ThCre:eLMO3 neurons was not different from control neurons (two-tailed paired t-test p = 0.2407, t = 1.459, df = 3) **(Figure 5G & I-J)**.

### ES fails to enhance recovery of sweating and anatomical sweat gland reinnervation

Every 2 weeks for 12 weeks after sciatic nerve transection and repair, the number of functional sweating droplets was counted on all 6 footpads of the hind paws to track the recovery of the sympathetic skin response **(Figure 7A)**, which is independent of sympathetic skeletal muscle activity (Delius et al., 1972; Derius, 1972; Hagbarth et al., 1972). The number of sweat droplets recovered at similar rates in the control and the ES groups over time, returning to about 73% of baseline at 12 weeks (2- way RM ANOVA, experimental group factor not significant) **(Figure 7B)**. This result is consistent with previous studies using the pilocarpine assay to track functional sympathetic recovery in a rodent model (Navarro and Kennedy, 1989). However, the most distal footpad, labeled as footpad 3 **(Figure 7C)**, only recovered to about 56% 12 weeks after injury (ordinary one-wave ANOVA, F = 10.09, p = 0.0006, post hoc Tukey’s multiple comparisons test: Intact vs Sham p = 0.0214, Intact vs ES p = 0.0008) **(Figure 7D-H)**. This most distal footpad was then chosen for anatomical reinnervation quantification as it was the footpad that exhibited the least sweating recovery **(Figure 7F-H)**.

ES did not enhance the sympathetic reinnervation of the most distal footpad 12 weeks after injury compared to Sham (ordinary one-way ANOVA, F = 13.72, p = 0.0002, post hoc Tukey’s multiple comparisons test: Intact vs Sham p = 0.0012, Intact vs ES p = 0.0008) **(Figure 8A & D)**. Both injury groups (Sham and ES) also exhibited a small degree of aberrant sweat gland reinnervation by YFP+ axons (Kruskal-Wallis test p = 0.0058, post hoc Dunn’s multiple comparisons test: Intact vs Sham p = 0.0588, Intact vs ES p = 0.0130) **(Figure 8B, E, & G)**. Consistent with our observation that sweating was not different between Sham and ES groups, total reinnervation of the sweat glands was similar between these groups (ordinary one-way ANOVA, F = 11.78, p = 0.0004, post hoc Tukey’s multiple comparisons test: Intact vs Sham p = 0.0034, Intact vs ES p = 0.0012) **(Figure 8C & F)**.

**Figure 7:**
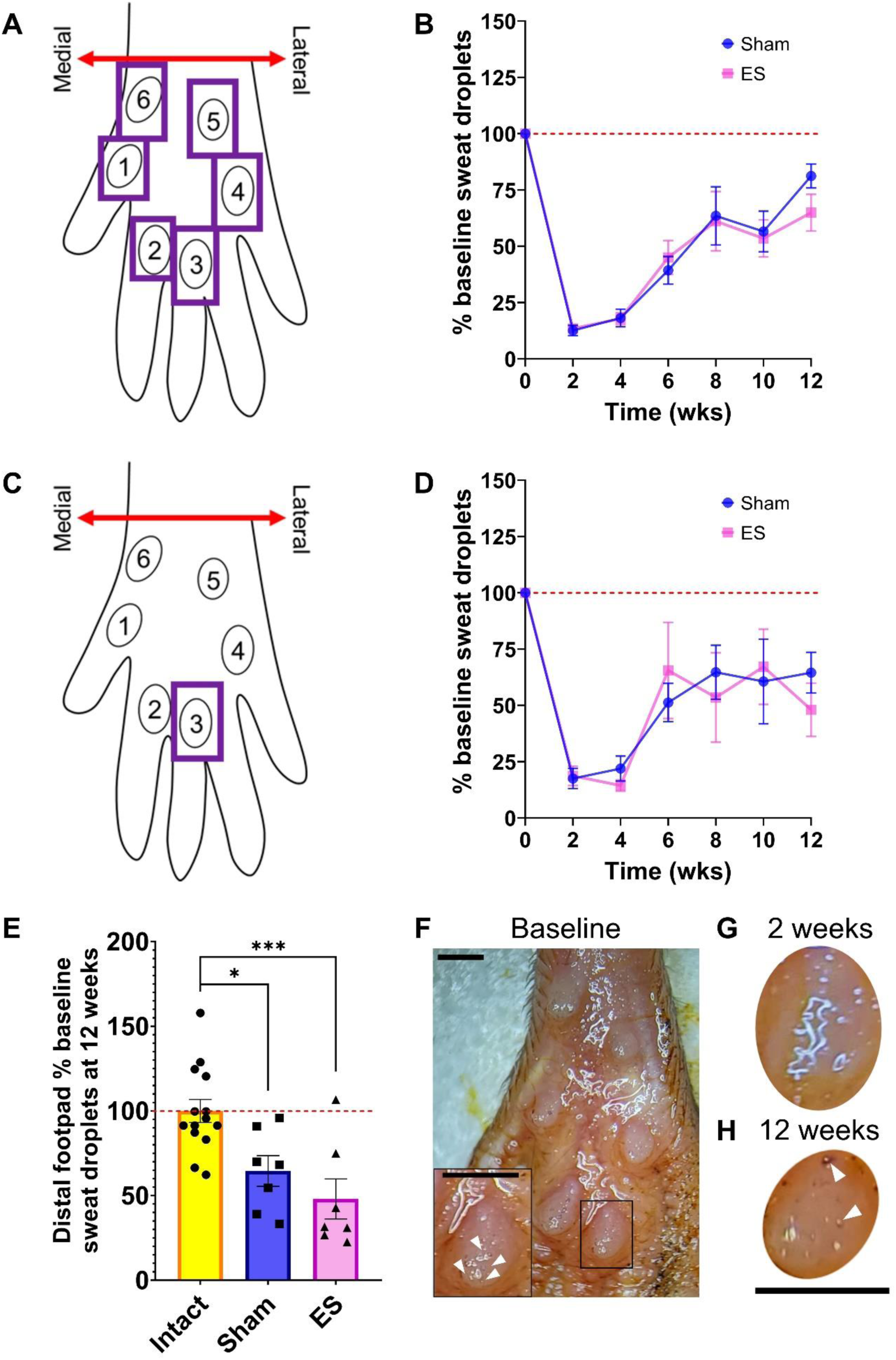
Electrical stimulation does not enhance the recovery of the sweating response with the greatest deficit observed at the most distal footpad. (A) Diagram of mouse hindlimb footpad orientation. Sweat droplets on all 6 footpads to evaluate total functional recovery. (B) Quantification of sweat droplets over time on all 6 footpads. (C) Diagram of mouse hindlimb footpad orientation, focusing on the most distal footpad labeled “3.” (D) Quantification of sweat droplets over time only on footpad 3. (E) Quantification of sweat droplets on footpad 3 compared to baseline 12 weeks after injury. One-way ANOVA with post hoc Tukey’s multiple comparisons test. * p < 0.05. *** p < 0.001. Data shown as mean ± SEM. (F) Representative images of footpad 3 with sweat droplets (arrows) at baseline, (G) 2 weeks after injury, and (H) 12 weeks after injury. Scale bars = 1 mm. n = 7 for Intact, Sham, and ES.

**Figure 8:**
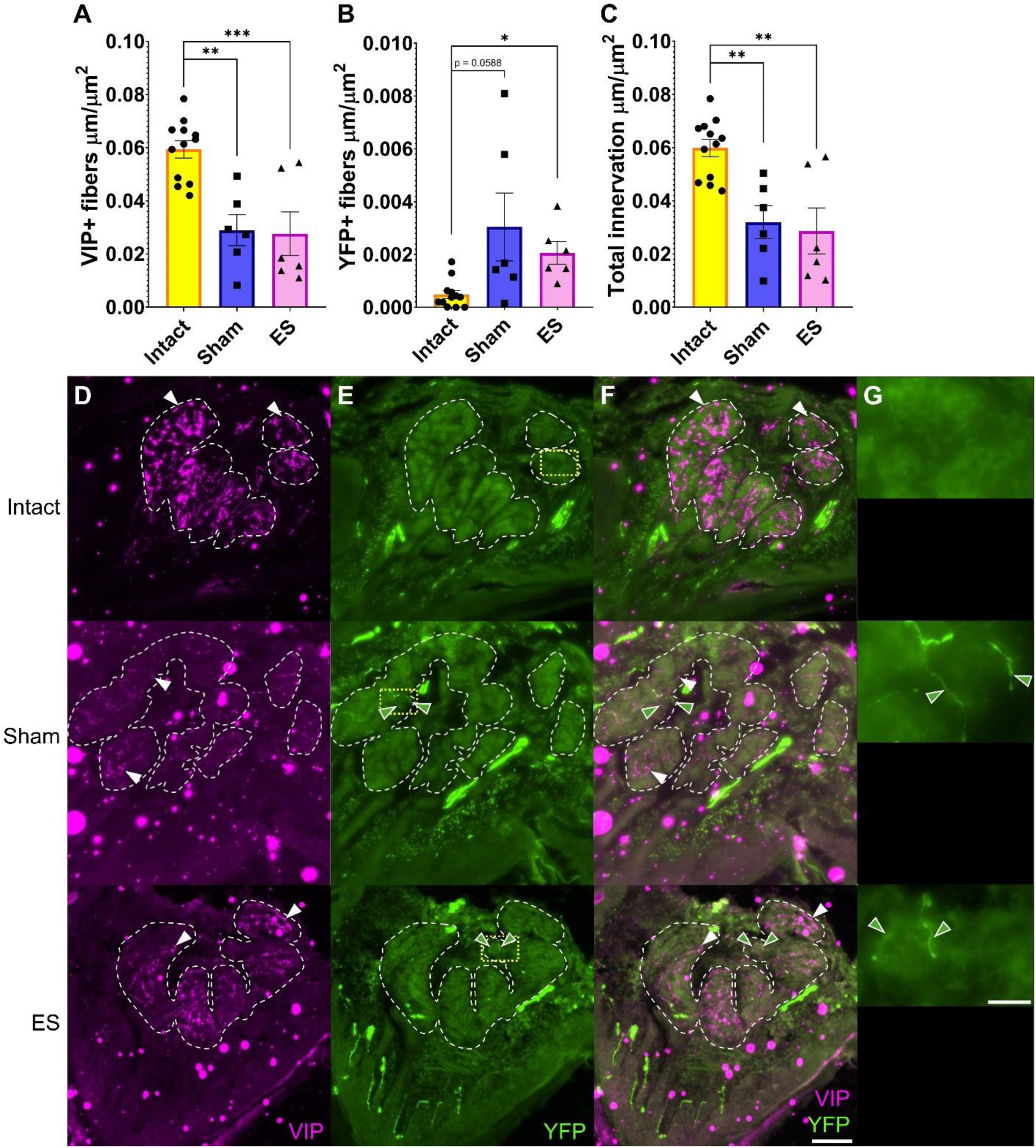
Reinnervation of the sweat glands in the most distal footpad was not improved by electrical stimulation, and aberrant reinnervation was observed in both injury groups. (A) Quantification of sympathetic fibers (VIP, vasoactive intestinal peptide, magenta), (B) aberrant innervation (YFP, yellow fluorescent protein, green), and (C) total innervation of the sweat gland area of the most distal footpad 12 weeks after sciatic nerve transection and repair. One-way ANOVA with post hoc Tukey’s multiple comparisons test. * p < 0.05. ** p < 0.01. *** p < 0.001. Data shown as mean ± SEM. Representative footpad sections with (D) sympathetic fibers (white arrows, Alexa Fluor® 647) in the sweat gland area (white outline), (E) aberrant reinnervation (green arrows, yellow outline, YFP) in the sweat gland area in the Sham and ES groups. (F) Merged. Scale bar = 100 μm. (G) Enlarged images of yellow outlined areas. Scale bar = 20 μm. n = 7 for Intact, Sham, and ES.

### ES enhances motor regeneration to the foot but not sympathetic regeneration

Fluorescent retrograde tracers were injected into the intrinsic foot muscles and the tibialis anterior (TA) 12 weeks bilaterally in unilaterally injured mice. Because most axons will have generated past 5 mm 12 weeks after injury, even in the untreated group, retrograde tracer was used to assess reinnervation of proximal muscle targets (TA) and distal muscle targets (intrinsic foot muscles). The spinal cord and lumbar sympathetic ganglia were collected 3 days after injection to allow time for the retrograde transport of the tracers.

Retrograde tracing from the intrinsic foot muscles 12 weeks post-PNI revealed that ES enhanced the number of motor neurons that have reinnervated these distal foot muscles compared to Sham (ordinary one-way ANOVA, F = 65.32, p < 0.0001, post hoc Tukey’s multiple comparisons test: Intact vs Sham p < 0.0001, Intact vs ES p < 0.0001, Sham vs ES p = 0.0443) **(Figure 9A)**. However, ES did not increase the number of postganglionic sympathetic neurons that had reinnervated the foot muscles compared to Sham (ordinary one-way ANOVA, F = 17.75, p < 0.0001, post hoc Tukey’s multiple comparisons test: Intact vs Sham p = 0.0004, Intact vs ES p < 0.0001, Sham vs ES p = 0.8567) **(Figure 10A)**. Neither injury groups were able to return to the baseline level of motor innervation of the intrinsic foot muscles 12 weeks after injury **(Figures 9A)**.

**Figure 9:**
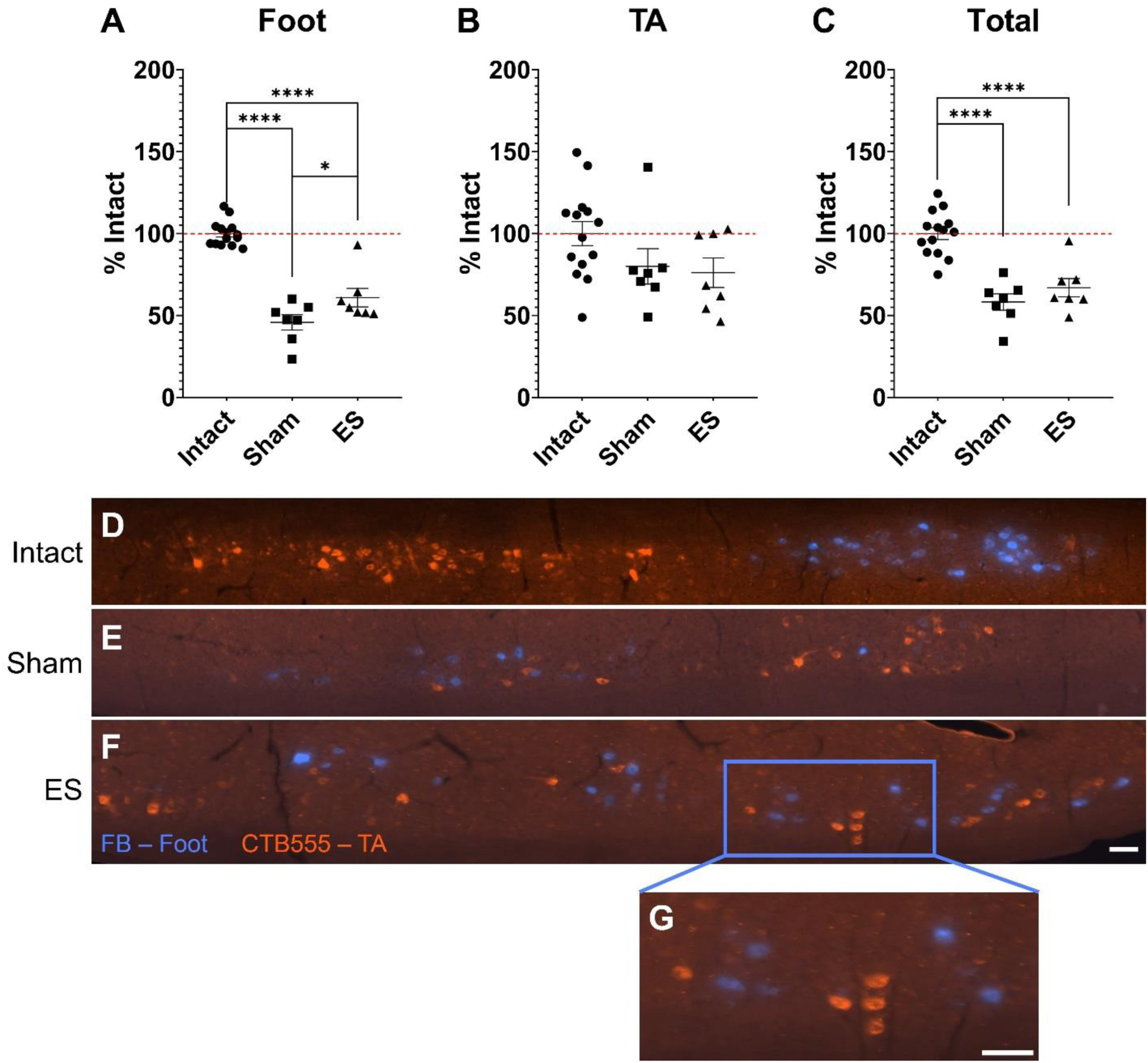
Electrical stimulation improves motor reinnervation of the foot while motor reinnervation of the tibialis anterior is complete 12 weeks after injury.Quantification of retrogradely-traced motor neurons innervating the (A) intrinsic foot muscles via fast blue (FB) and (B) tibialis anterior (TA) via Cholera Toxin Subunit B Alexa Fluor™ 555 (CTB555) compared to the intact side. (C) Total number of retrogradely-labeled motor neurons from the intrinsic foot muscles and TA compared to the intact side. One-way ANOVA with post hoc Tukey’s multiple comparisons test. * p <0.05. **** p < 0.0001. Data shown as mean ± SEM. Representative images of the intrinsic foot muscles (blue) and TA (red) motor pools in the spinal cord from (D) uninjured (Intact), (E) Sham stimulated, and (F) electrically-stimulated (ES) mice. (G) Enlarged image of labeled motor neurons. Scale bars = 500 μm. n = 7 for Intact, Sham, and ES.

**Figure 10:**
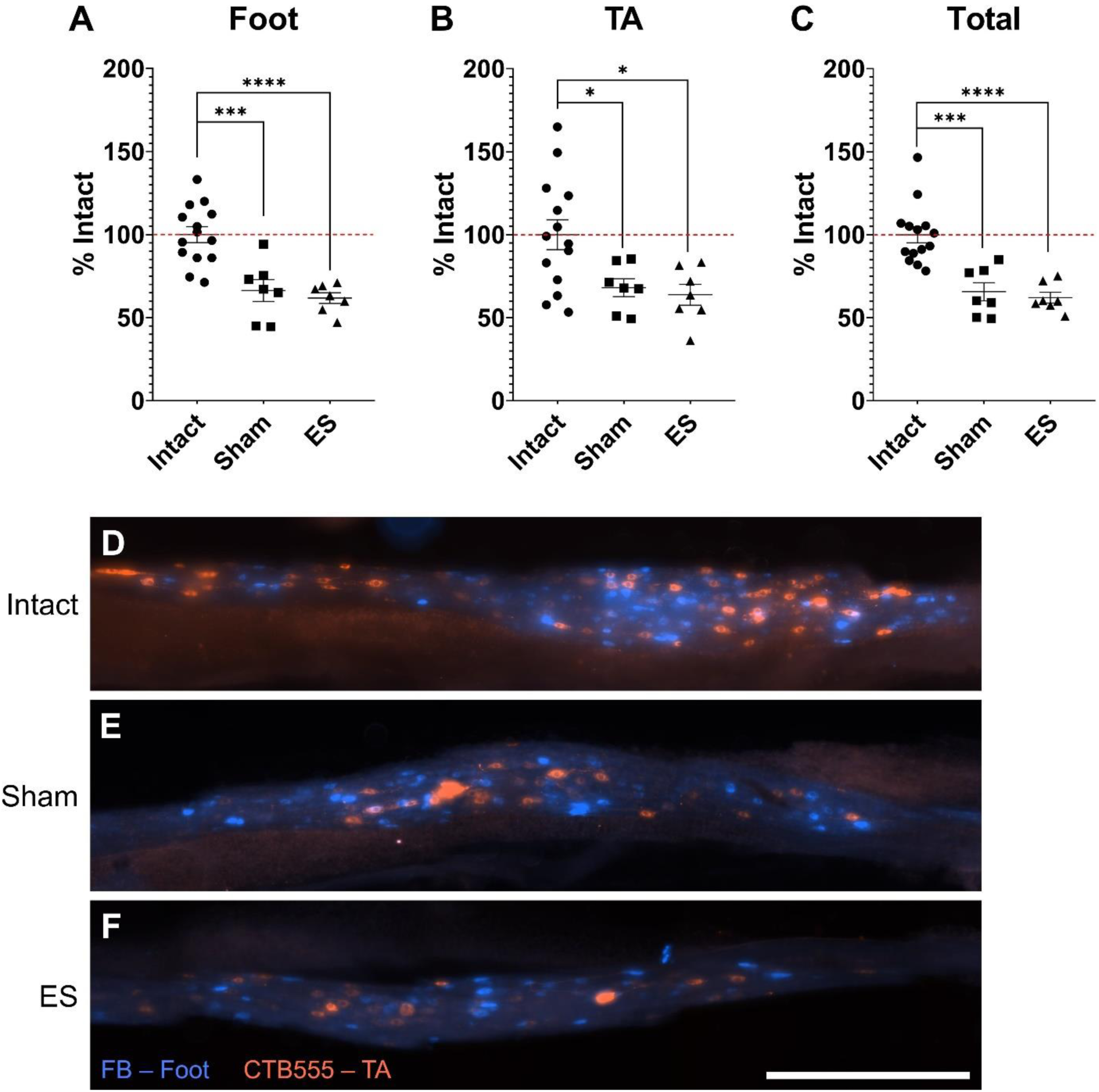
Sympathetic reinnervation of the intrinsic foot muscles and tibialis anterior is not enhanced by electrical stimulation. Quantification of retrogradely- traced lumbar sympathetic neurons innervating the (A) intrinsic foot muscles fast blue (FB) and (B) tibialis anterior (TA) via Cholera Toxin Subunit B Alexa Fluor™ 555 (CTB555) compared to the intact side. (C) Total number of retrogradely-labeled sympathetic neurons from the intrinsic foot muscles and TA compared to the intact side. One-way ANOVA with post hoc Tukey’s multiple comparisons test. * p < 0.05. Data shown as mean ± SEM. Representative images of the labeled neurons that innervate the intrinsic foot muscles (blue) and TA (red) in L2 sympathetic ganglion from (D) uninjured (Intact), (E) Sham stimulated, and (F) electrically-stimulated (ES) mice. Scale bar = 500 μm. n = 7 for Intact, Sham, and ES.

### Motor reinnervation of the tibialis anterior is complete by 12 weeks after injury, but sympathetic reinnervation remains incomplete

On the other hand, the TA, the more proximal target, exhibited near complete reinnervation by motor neurons 12 weeks post-PNI **(Figure 9B)**. The number of retrogradely-traced neurons in both injury groups was no different than that exhibited in Intact muscle. On the other hand, sympathetic reinnervation of the TA remained deficient (ordinary one-way ANOVA, F = 5.791, p = 0.0086, post hoc Tukey’s multiple comparisons test: Intact vs Sham p = 0.0399, Intact vs ES p = 0.0184, Sham vs ES p = 0.9521) **(Figure 10B)**.

### ES does not remedy changes in skeletal muscle cellular energy charge after PNI and fails to reverse muscle atrophy

Twelve weeks after PNI, the gastrocnemius (GA) was collected bilaterally from unilaterally injured mice and processed for purine quantification via HPLC to find energy charge, which is a sensitive indicator of cellular energy status and energy consumption (Atkinson, 1968, 1971; Friess et al., 2002; Metallo and Vander Heiden, 2013). The ATP, ADP, and AMP values were normalized to total protein content of the sample prior to calculating the energy charge.

There was no significant difference in the energy charge between Intact and Sham, a difference was found between Intact and ES (ordinary one-way ANOVA, F = 5.316, p = 0.0119, post hoc Tukey’s multiple comparisons test: Intact vs ES p = 0.0140) **(Figure 11A)**. Upon determining that the sample size was insufficient to reveal the differences between groups, estimation statistics were used to focus on the effect size (Ho et al., 2019). Both Sham and ES exhibited a decrease in energy charge compared to Intact when using estimation statistics (p = 0.0302 and p = 0.0026, respectively) (Figure 11B).

**Figure 11:**
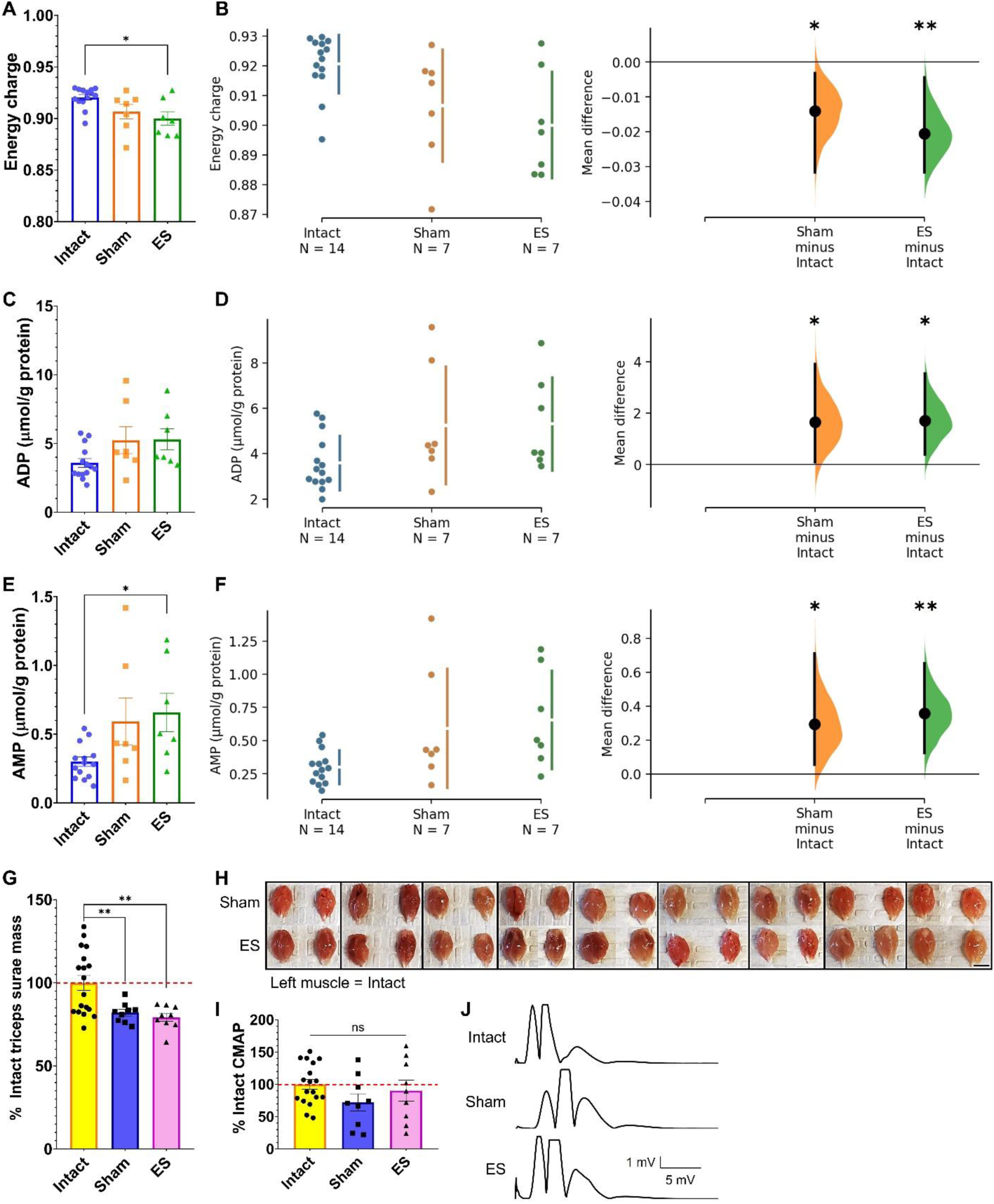
Electrical stimulation does not improve deficits in skeletal muscle cellular energy charge after peripheral nerve injury. (A, C, E) Energy charge,normalized ADP, and normalized AMP in the lateral head of the gastrocnemius. Ordinary one-way ANOVA with post hoc Tukey’s multiple comparisons test or Kruskal-Wallis test with post hoc Dunn’s multiple comparisons test. (B, D, F) Cumming estimation plot with the raw data on the left and the mean differences, depicted as singular dot, on the right plotted as bootstrap sampling distributions with 95% confidence intervals. n = 7 for Intact, Sham, and ES. (G) Triceps surae muscle weights represented as a percentage of the intact side. One-way ANOVA with post hoc Tukey’s multiple comparisons test. * p< 0.05. ** p < 0.01. (H) All muscles included in (G) shown in pairs with the intact triceps surae in each pair on the left. Scale bar = 5 mm. (I) Maximum compound muscle action potential (CMAP) from the gastrocnemius as a percentage of the contralateral intact side. (K) Representative rectified CMAP traces. Data shown as mean ± SEM. n = 9 for Intact, Sham, and ES.

There was also no difference in ADP levels between any of the groups (Kruskal-Wallis test, p = 0.0568) **(Figure 11C)**, but when estimation statistics were employed, ADP levels were elevated in both injury groups (Intact vs Sham p = 0.0492, Intact vs ES p = 0.022) **(Figure 11D)**. AMP values were also elevated after injury, with a significant elevation in the ES group over Intact (Kruskal-Wallis test p = 0.0303, post hoc Dunn’s multiple comparisons test: Intact vs ES p = 0.0359) **(Figure 11E)**. Estimation statistics revealed that *both* injury groups exhibited elevated AMP compared to the uninjured group (Intact vs Sham p = 0.0212, Intact vs ES p = 0.0028) **(Figure 11F)**. Finally, ES failed to reverse the muscle atrophy observed after chronic PNI (Brown-Forsythe

ANOVA test, F (DFn, DFd) = 13.68 (2.000, 27.36), p < 0.0001, post hoc Tamhane’s T2 multiple comparisons test: Intact vs Sham p = 0.0041, Intact vs ES p = 0.0014) **(Figure 11G-H)**; however, the amplitudes of the maximum CMAPs elicited in the gastrocnemius had returned to baseline intact levels in both injury groups (ordinary one-way ANOVA, F = 1.597, p = 0.2177) **(Figure 11I-J)**

### ES rescues acetylcholine receptor content at the NMJ but not β2 adrenergic receptors

The presence of acetylcholine receptors (AChRs) and ADRB2s at NMJs in the GA was visualized via IHC 12 weeks after injury. Using fluorescence intensity as an indicator of protein content, ES was found to partially rescue the deficit in AChR content at the NMJ compared to Sham although not back to Intact levels (Brown-Forsythe ANOVA test, F (DFn, DFd) = 44.41 (2.000, 642.1), p < 0.0001, post hoc Games-Howell’s multiple comparisons test: Intact vs Sham p < 0.0001, Intact vs ES p = 0.0010, Sham vs ES p < 0.0001) **(Figure 12A & D)**. Furthermore, ES increased the NMJ area (Brown-Forsythe ANOVA test F (DFn, DFd) = 8.909 (2.000, 595.5), p = 0.0002, post hoc Games-Howell’s multiple comparisons test: Intact vs Sham p = 0.0908, Intact vs ES p = 0.0393, Sham vs ES p = 0.0002) **(Figure 12B)**.

**Figure 12:**
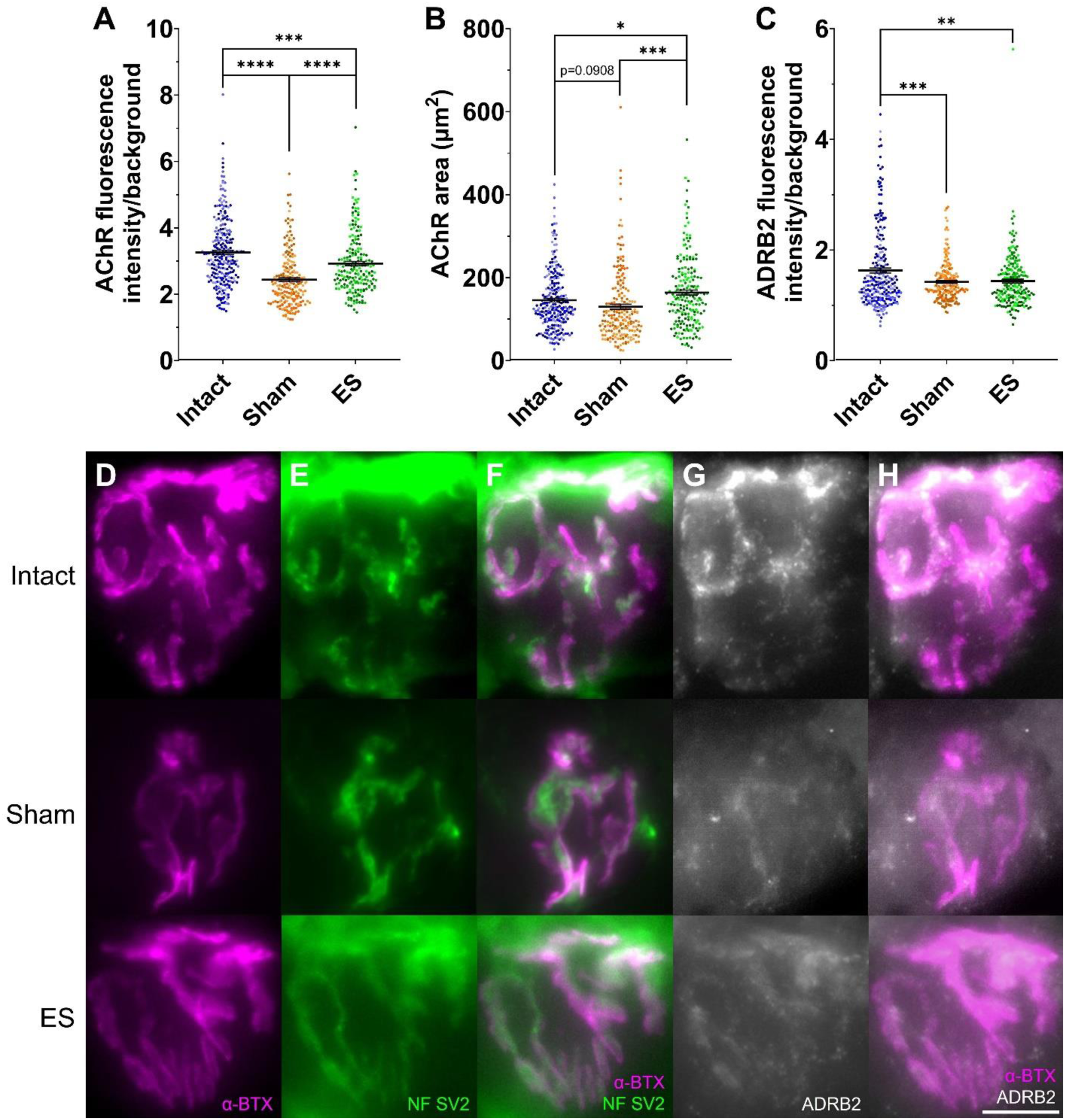
Electrical stimulation partially rescues acetylcholine receptor content at the neuromuscular junction but fails to improve β2-adrenergic receptor content. (A) Acetylcholine receptor (AChR) fluorescence quantification at the neuromuscular junction (NMJ). (B) Area occupied by AChRs. (C) β2-adrenergic receptor (ADRB2) fluorescence quantification at the NMJ. Representative images of Intact, Sham, and ES NMJs showing the following: (D) AChR labeled by α-bungarotoxin (α- BTX Alexa Fluor® 488), (E) motor reinnervation via neurofilament-H (NF) and synaptic vesicle protein 2 (SV2) (Alexa Fluor® 647), (F) AChR overlaid by NF and SV2, (G) ADRB2 (Alexa Fluor® 555), and (H) AChR overlaid by ADRB2. Pseudocolored for ease of visualization. Intact n = 5, Sham n = 4, ES n = 4.

In contrast, despite the return of motor reinnervation **(Figure 12E-F)**, ES was unable to rescue the deficits in ADRB2 content at the NMJ (Brown-Forsythe ANOVA test F (DFn, DFd) = 10.73 (2.000, 561.6), p < 0.0001, Intact vs Sham p = 0.0002, Intact vs ES p = 0.0025, Sham vs ES p = 0.9353) **(Figure 12C & G-H)**. This is consistent with the deficits seen in sympathetic regeneration to the more proximal muscles of the hindlimb (TA) in this same cohort of mice **(Figure 10B)**.

## DISCUSSION

The sympathetic nervous system is crucial for a variety of bodily homeostatic mechanisms, including but not limited to, sweating and skeletal muscle mitochondria metabolism (Robinson et al., 2011; Khan et al., 2016; Hu et al., 2018; Azevedo Voltarelli et al., 2021); therefore, promoting the regeneration of postganglionic sympathetic axons is important. In our studies, neither ES nor a CL enhanced the acute sympathetic regeneration after sciatic nerve transection and repair. However, both of these interventions successfully enhanced acute motor and sensory regeneration, consistent with previous research (McQuarrie et al., 1977; Forman et al., 1981; McQuarrie, 1986; Al-Majed et al., 2000; Brushart et al., 2005; Geremia et al., 2007; Gordon et al., 2009; Hoffman, 2010; Wong et al., 2015). Furthermore, the use of BL-OG to selectively target postganglionic sympathetic activity inhibited their growth. These findings demonstrate that the postganglionic lumbar sympathetic neurons exhibit a distinct response to neuronal stimulation compared to their motor and sensory counterparts.

ES recruits axons into activity from the largest myelinated axons to the smallest unmyelinated axons (Blair and Erlanger, 1933; Fang and Mortimer, 1991; Singh et al., 2000; Llewellyn et al., 2010). Therefore, the clinically relevant ES paradigm is unlikely to recruit small, unmyelinated postganglionic sympathetic fibers into activity. The CL paradigm, which relies on a nerve crush before a test lesion to stimulate regeneration, does not depend on the electrical properties of the axons and induces the upregulation of transcription factors, such as ATF3 (Sachs et al., 2010). While McQuarrie et al. found that the sympathetic axons in the sciatic nerve are inhibited by a CL, others found that sympathetic regeneration is enhanced with a CL (McQuarrie et al., 1978; Navarro and Kennedy, 1990a; Shoemaker et al., 2005). Notably, one of the studies showing enhanced sympathetic regeneration studied the superior cervical ganglia, and different anatomical levels of the sympathetic ganglia exhibit different molecular signatures (Furlan et al., 2016); therefore, lumbar sympathetic neurons likely react differently to a CL compared to cervical sympathetic neurons. Despite the ability of a CL to induce changes in all axons in the sciatic nerve through prior injury, it was also unable to enhance sympathetic regeneration in our experiments.

To isolate the activity of postganglionic sympathetic neurons, BL-OG was employed using a ThCre:eLMO3 mouse, which expresses an excitatory luminopsin eLMO3 in TH+ neurons. The luminopsin is a fusion of a light-generating luciferase and a channelrhodopsin. When the luciferase substrate CTZ is introduced, light is generated and causes neuronal excitation **(Figure 3A)** (Berglund et al., 2016; Gomez-Ramirez et al., 2020; Berglund et al., 2021). Due to the nonspecific nature of ES and CL as well as the potential inability of ES to stimulate small-caliber sympathetic axons, as supported by the lack of differential gene expression in lumbar sympathetic neurons in response to ES, with only 52 upregulated genes with the addition of ES in the context of nerve injury vs 223 upregulated genes from nerve injury alone **(See Data Availability)**, BL-OG was employed as a novel technique to target sympathetic activity.

Interestingly, BL-OG inhibited sympathetic regeneration as evidenced by fluorescent retrograde tracing 2 weeks after injury **(Figure 5D-F)**. The decrease in the number of retrograde-traced sympathetic neurons in the BL-OG paradigm was not replicated in the ES and CL paradigms **(Figure 2I-L)**, indicating that specific targeting of sympathetic activity has an additional inhibitory effect. Furthermore, BL-OG reduced the branching of sympathetic neurons *in vitro* **(Figure 5H)**. Elongation, represented by the enclosing radius, and branching, represented by the branching index, are both important components of regeneration (Caroni, 1997; Xu et al., 2008; Lemaitre, 2021).

Although excitotoxicity may be a concern due to the long-lasting bioluminescence **(Figure 3C)** and sustained negative deflection in the LFPs **(Figure 4E)**, this BL-OG model has been used previously, *in vivo* and *in vitro*, with the same CTZ doses and has successfully enhanced motor and sensory regeneration in PNI and spinal cord injury models (Crespo et al., 2021; English et al., 2021; Medendorp et al., 2021; Ecanow et al., 2022; Ikefuama et al., 2022). The results of the ES and CL acute experiments, however, reassure us that the congruous results in the BL-OG model further support that neuronal stimulation does not enhance sympathetic regeneration. Future mechanistic studies are necessary to understand how to combat the block in sympathetic regeneration after injury.

Upon investigation of the long-term effects of ES on sympathetic regeneration, ES again failed to enhance both functional and anatomical sympathetic regeneration. Neither the sweating response nor the reinnervation of the distal sweat glands were significantly enhanced over Sham. Furthermore, both injury groups exhibited a small degree of aberrant sweat gland reinnervation evidenced by YFP+ axons in the sweat gland area **(Figure 8B, E, & G)**. Injured neurons undergo neural plasticity after PNI, switching from a transmitter state to a regenerative state while expressing molecules not normally expressed in adult neurons (Navarro et al., 2007). In the YFP-16 mice used in this study, YFP expression is found in medium- and large-diameter sensory neurons all the way to their peripheral terminals such as Meissner corpuscles and muscle spindles but not in small-diameter nociceptive neurons (Taylor-Clark et al., 2015). Therefore, the YFP+ axons in the sweat gland area 12 weeks after injury were likely medium- or large-diameter sensory neurons.

After injury, there is a marked increase in neuropeptide Y (NPY), which is typically barely detectable, in medium and large sensory neurons (Wakisaka et al., 1991; Ohara et al., 1994). NPY is present in sympathetic precursors and is co-expressed in norepinephrine-positive sympathetic fibers primarily in blood vessels and vascular beds (Lundberg et al., 1982; Lundberg et al., 1983; Ekblad et al., 1984; Lundberg et al., 1984; Tyrrell and Landis, 1994; Hall and MacPhedran, 1995). NPY+ fibers have also been found around eccrine sweat glands (Björklund et al., 1986; Johansson, 1986; Wallengren et al., 1987). Although vasoactive intestinal peptide has been shown to be upregulated in sensory neurons after nerve injury, this molecular change is found exclusively in small sensory neurons (Shehab et al., 1986), which may not express YFP in YFP-16 mice (Taylor-Clark et al., 2015). Hence, molecular changes exhibited in medium and large sensory axons after injury make them histologically similar to postganglionic sympathetic axons. Whether the infiltration of YFP+ axons into previously unoccupied areas before injury represents a true phenotypic switch requires further investigation.

Muscle fluorescent retrograde tracing revealed that 12 weeks after injury, the number of motor neurons that had reinnervated more proximal muscles, such as the TA, had returned to nearly baseline levels **(Figure 9B)**. Despite the near complete return of motor reinnervation to the TA, sympathetic reinnervation of the TA remained deficient (Figure 10B).

Long-term denervation has been linked to an overall decrease in AChR (Rochkind and Shainberg, 2017). In the GA, ES was able to partially rescue the AChR content at the NMJ **(Figure 12A & D)**. This is consistent with previous work that showed that AChR content increases with direct ES of muscle cells and with chronic low-frequency (10 Hz) stimulation through an intact nerve (O’Reilly et al., 2003; Lozano et al., 2016), but to our knowledge, this is the first report of increased AChR in response to ES of an injured nerve. ADRB2 content, on the other hand, was decreased and unaffected by ES **(Figure 12C & G-H)**, supporting the lack of effect on sympathetic regeneration.

Perturbations in mitochondrial function have been previously documented in denervated skeletal muscle, and more recently, sympathetic signaling in skeletal muscle has been implicated in mitochondrial function (Khan et al., 2016; Yang et al., 2020).

Here, we observed a modest, but significant, persistent reduction of energy charge in the GA due to ADP and AMP elevations after PNI that were not remedied by ES **(Figure 11A-F)**, though all energy charge values remained in the normal range of 0.8-0.95 for eukaryotic cells (Atkinson, 1968). These findings indicate that there is an increased pool of low-energy molecules in skeletal muscle cells, reminiscent of changes in the energy charge in human skeletal muscle after exercise-induced exhaustion (Norman et al., 1987). By 12 weeks after injury, CMAP amplitudes in the GA had returned to intact values **(Figure 11I-J)**. Therefore, the persistent deficits exhibited in the energy charge 12 weeks after injury can likely be attributed to the lack of sympathetic reinnervation,

rather than the lack of motor reinnervation, in the muscle.

ES applied directly to the nerve for nerve regeneration has not been shown to improve the muscle atrophy associated with denervation and reinnervation, unlike direct muscle ES (Gibson et al., 1988; Hasegawa et al., 2011; Hirose et al., 2013; Willand et al., 2013). Changes in mitochondrial function are linked to muscle atrophy (Yang et al., 2020; Chen et al., 2023), and ADRB2 agonists have been shown to increase muscle mass in normal muscle and reduce muscle wasting in dystrophic muscle in animal models (Maltin et al., 1987; Kim and Sainz, 1992; Zeman et al., 1994). Furthermore, inhibition of the ubiquitin proteasome system and the expression of muscle atrophy-related genes *Atrogen-1* and *MuRF1 in vitro* by norepinephrine implies the antiproteolytic role of sympathetic signaling in skeletal muscle (Silveira et al., 2014).

Because the GA has likely attained complete motor reinnervation at 12 weeks post-injury but incomplete sympathetic reinnervation, the inability of ES to rescue atrophy is likely due, in part, to the lack of sympathetic reinnervation and subsequent lack of postsynaptic ADRB2 in the context of chronic PNI.

## CONCLUSIONS

The sympathetic nervous system modulates the activity of nearly every structure in the human body and plays a critical role in maintaining bodily homeostasis. The regeneration of postganglionic sympathetic axons in response to the clinically relevant ES paradigm, however, has not been well-studied. In this series of experiments, we found that ES is unable to enhance the regeneration of these small-caliber unmyelinated postganglionic sympathetic axons, unlike their motor and sensory counterparts.

This inability of sympathetic axons to regenerate in response to ES may lead to persistent changes in skeletal muscle cellular energy status and muscle bulk, which will require more in-depth investigations into the underlying mechanisms. Although the negligible effects of ES on sympathetic regeneration could be due to the inability of the ES paradigm to recruit small-caliber unmyelinated axons, both the CL and BL-OG also produced negligible or negative effects. Therefore, the molecular signature governing sympathetic regeneration is likely to be vastly different than that governing sensory and motor regeneration. This distinct molecular signature needs to be further investigated to develop therapeutic options that will enhance the regeneration of all axons.

## ACKNOWLEDGEMENTS

This work was supported by the National Institutes of Health, National Institute of Neurological Disorders and Stroke award number K01NS124912 (PJW) and National Institute of General Medical Sciences award number R01GM132598 (HCH) and in part by a developmental grant from the NIH-funded Emory Specialized Center of Research Excellence in Sex Differences U54AG062334 (PJW).

Support was also provided by the Emory University Integrated Cellular Imaging Core Facility (RRID:SCR_023534) and the Emory HPLC Bioanalytical Core (EHBC). The EHBC is subsidized by the Emory University School of Medicine and is one of the Emory Integrated Core Facilities. The content is solely the responsibility of the authors and does not necessarily reflect the official views of the National Institutes of Health. I would like to thank Dr. Ken Berglund for his guidance on the BL-OG system, Dr. Shawn Hochman and Dr. Celia (Yaqing) Li for their advice and expertise on sympathetic activity and Dr. Victor Faundez for his feedback on the interpretation of muscle purine content and expertise on mitochondrial function. Thank you to my thesis committee members (Dr. Hochman, Dr. Faundez, Dr. Arthur English, and Dr. Nicholas Boulis) for their contributions and input on this project. Additionally, I would like to thank HaoMin SiMa, research specialist, for 3D-printing a phone mount for our stereo microscope that allowed for the capturing of some of the images presented in this article. The 3D model was made available by OpenOcular and can be accessed via this link: www.thingiverse.com /thing:5186470. Thank you to Kevin Patel, undergraduate student, and Kate Pollack, research specialist, for assisting in some of the surgical procedures. Finally, thank you to Dr. Jingsheng Gu for his diligent maintenance of the mouse colony.

## DATA AVAILABILITY

The datasets generated during and/or analyzed during the current study are available from the corresponding author on reasonable request.

---

**Supplemental Figure 1:**
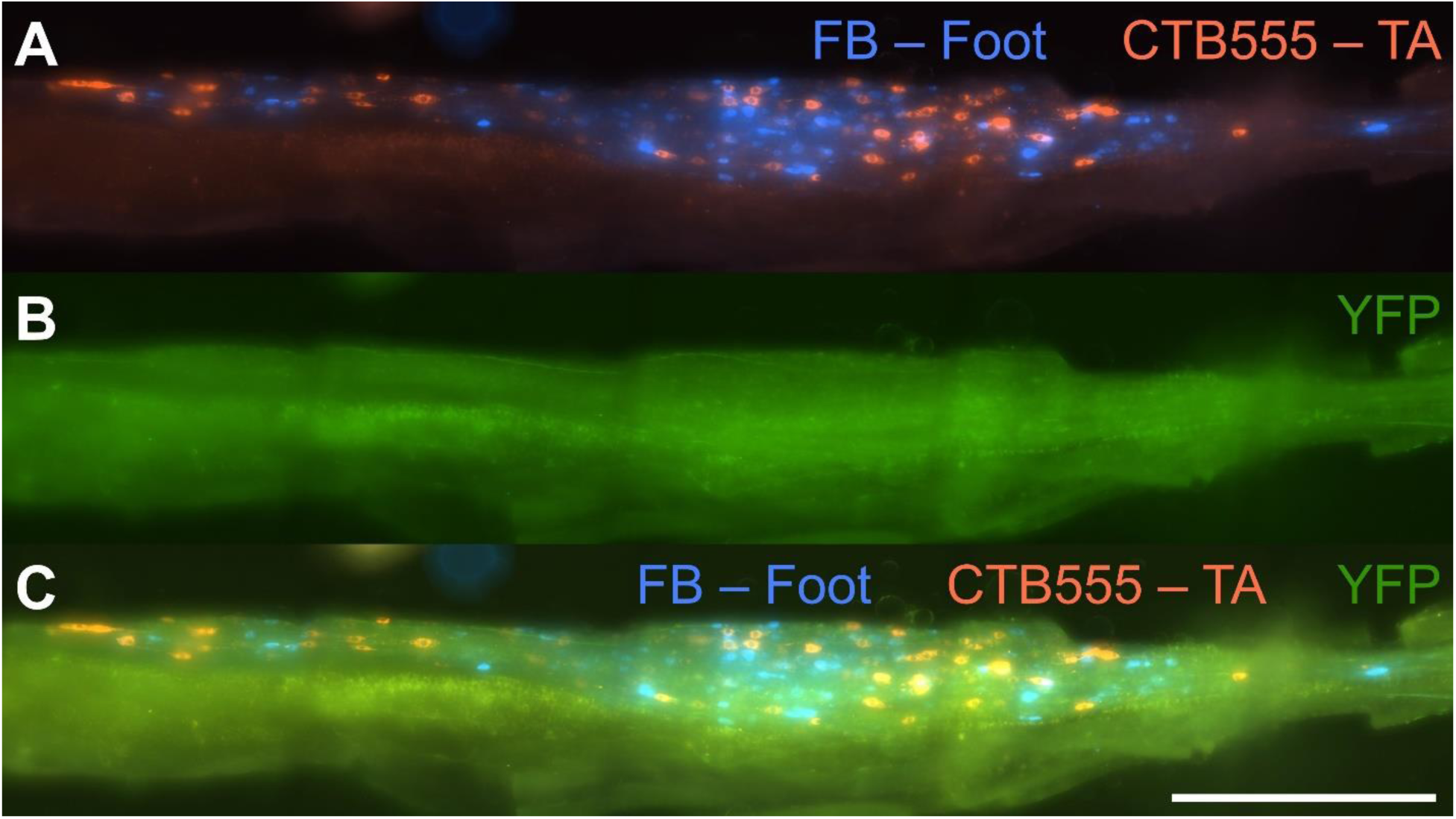
A YFP-16 mouse exhibits negligible expression of yellow fluorescent protein (YFP) in postganglionic sympathetic neurons. (A) An L2 lumbar sympathetic ganglion with blue and red neurons from the intrinsic foot muscles and tibialis anterior, respectively. (B) Visualization of YFP on the FITC channel. (C) Merged. Scale bar = 500 μm.

**Supplemental Figure 2:**
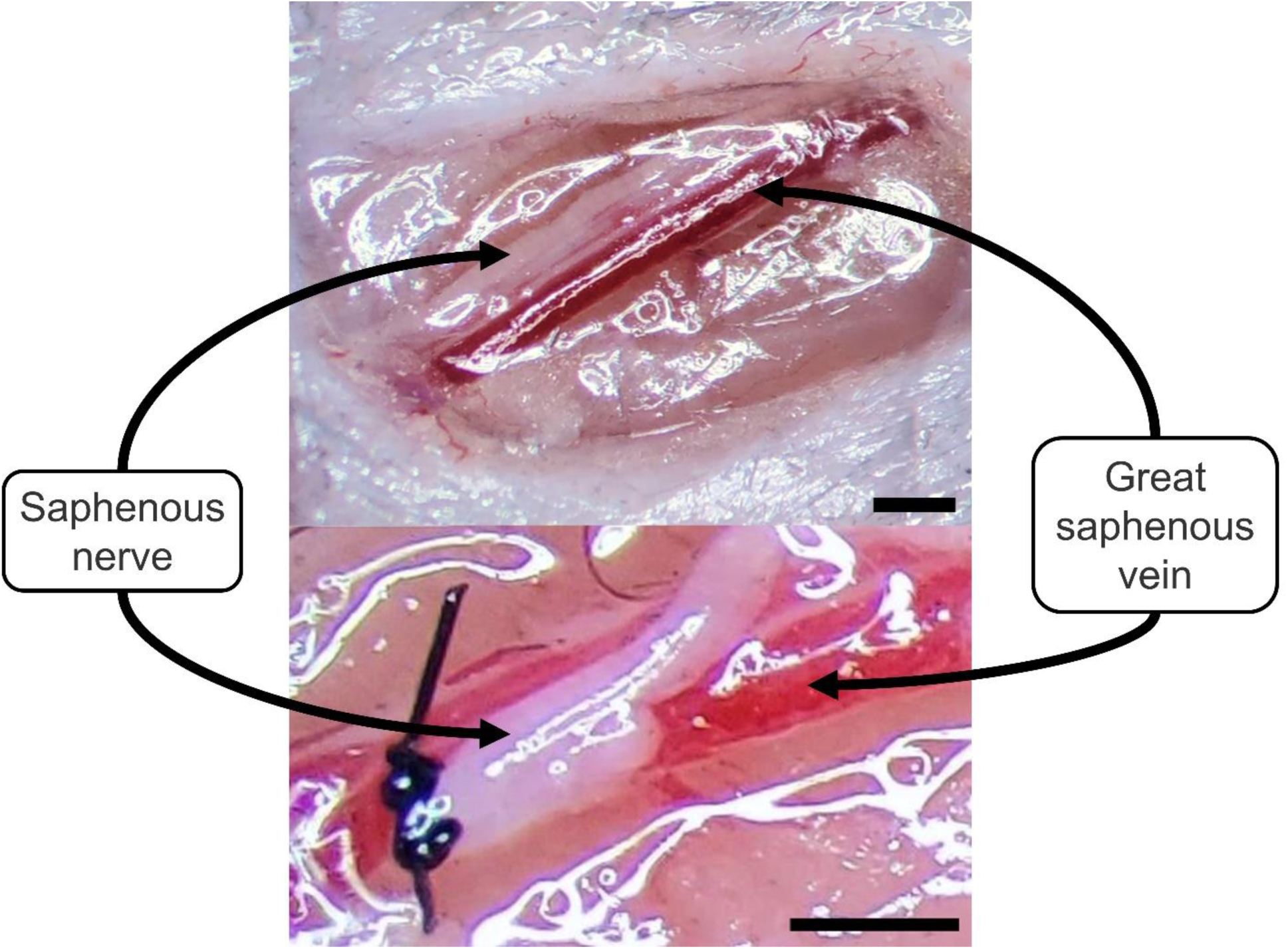
Identification (upper panel) and ligation (lower panel) of the saphenous nerve with 8-0 nylon suture near the great saphenous vein. Scale bars = 0.5 mm.

